# Cryo-EM of a Divergent Herpesvirus Reveals Structural Conservation and Novelty Including a Portal-Vertex Tegument Protein with Multiple Macrodomain-like Folds

**DOI:** 10.64898/2026.02.13.705735

**Authors:** Amit Meir Ben Efraim, Filip Petrov, Marion McElwee, James Streetley, Liisa Holm, Joe Grove, Andrew J. Davison, Frazer J. Rixon, David Bhella

**Affiliations:** MRC – University of Glasgow Centre for Virus Research, Sir Michael Stoker Building, Garscube Campus, 464 Bearsden Road, Glasgow, G61 1QH, United Kingdom

**Author notes:** Organismal and Evolutionary Biology Research Program, Faculty of Biological and Environmental Sciences & Institute of Biotechnology, Helsinki Institute of Life Sciences, University of Helsinki, Helsinki, Finland (Liisa Holm). Authors contributed equally (Amit Meir Ben Efraim, Filip Petrov).

## Abstract

Ictalurid herpesvirus 1 (channel catfish virus) is an evolutionarily distant relative of human herpesviruses, from which it is thought to have diverged >400M years ago. Using cryogenic electron microscopy (cryo-EM) combined with symmetry-breaking and particle subtraction approaches, we determined structures of both the immature capsid and virion of IcHV-1. Due to limited genome annotation, we used the machine learning-based tool *ModelAngelo* for *de novo* model building, enabling unambiguous protein identification even at marginal resolutions. Notably, the IcHV-1 virion was found to have a substantial and elaborate portal-vertex associated tegument (PVAT) complex. Overall, we determined the identities and structures of ten IcHV-1 proteins: the major capsid protein; the triplex proteins; two novel virion-associated inner tegument proteins; the portal protein; and a further four PVAT proteins. Our findings reveal a high degree of fold conservation in the core capsid proteins when compared with those of human herpesviruses, but also considerable structural novelty, including for the first time in a herpesvirus, identification of a protein that has four putative macrodomains.

## Introduction

Herpesviruses are large DNA viruses that establish latency in their hosts, causing life-long infection. They include many notable human pathogens including varicella-zoster virus (VZV) – the cause of chicken pox and herpes zoster (shingles); Epstein Barr virus (EBV) – the cause of infectious mononucleosis; human cytomegalovirus (HCMV) – the leading viral cause of congenital abnormalities; and herpes simplex viruses (HSV) 1 and 2 that cause cold-sores and genital lesions respectively, and sometimes sight-threatening keratitis, or severe encephalitis. The order *Herpesvirales* contains three virus families – the *Orthoherpesviridae* which includes viruses of mammals (including humans), birds and reptiles, the *Malacoherpesviridae* which includes viruses of molluscs and the *Alloherpesviridae* which includes viruses of amphibia and fish (*1*). Herpesviruses are highly host-specific and are thought to have evolved primarily by co-speciation with their hosts (*2*). Thus, the divergence between the herpesvirus families likely follows that of their hosts, with O*rthoherpesviridae* and *Alloherpesviridae* having diverged more than 400 million years ago (*3*). Evidence of a much older common ancestor between herpesviruses and the *Caudoviricetes*, which are tailed bacteriophages that infect both prokaryotes and archaea, is found in their common capsid assembly pathways and the presence of the HK97/Johnson fold in their major capsid proteins (*4*) (figure 1A). Since conservation of protein folds has revealed unsuspected and surprising evolutionary relationships between viruses (*5*), structure analysis of evolutionarily distant herpesviruses has the potential to further inform our understanding of herpesvirus evolution and highlight features specific to the host species.

**Figure 1.**
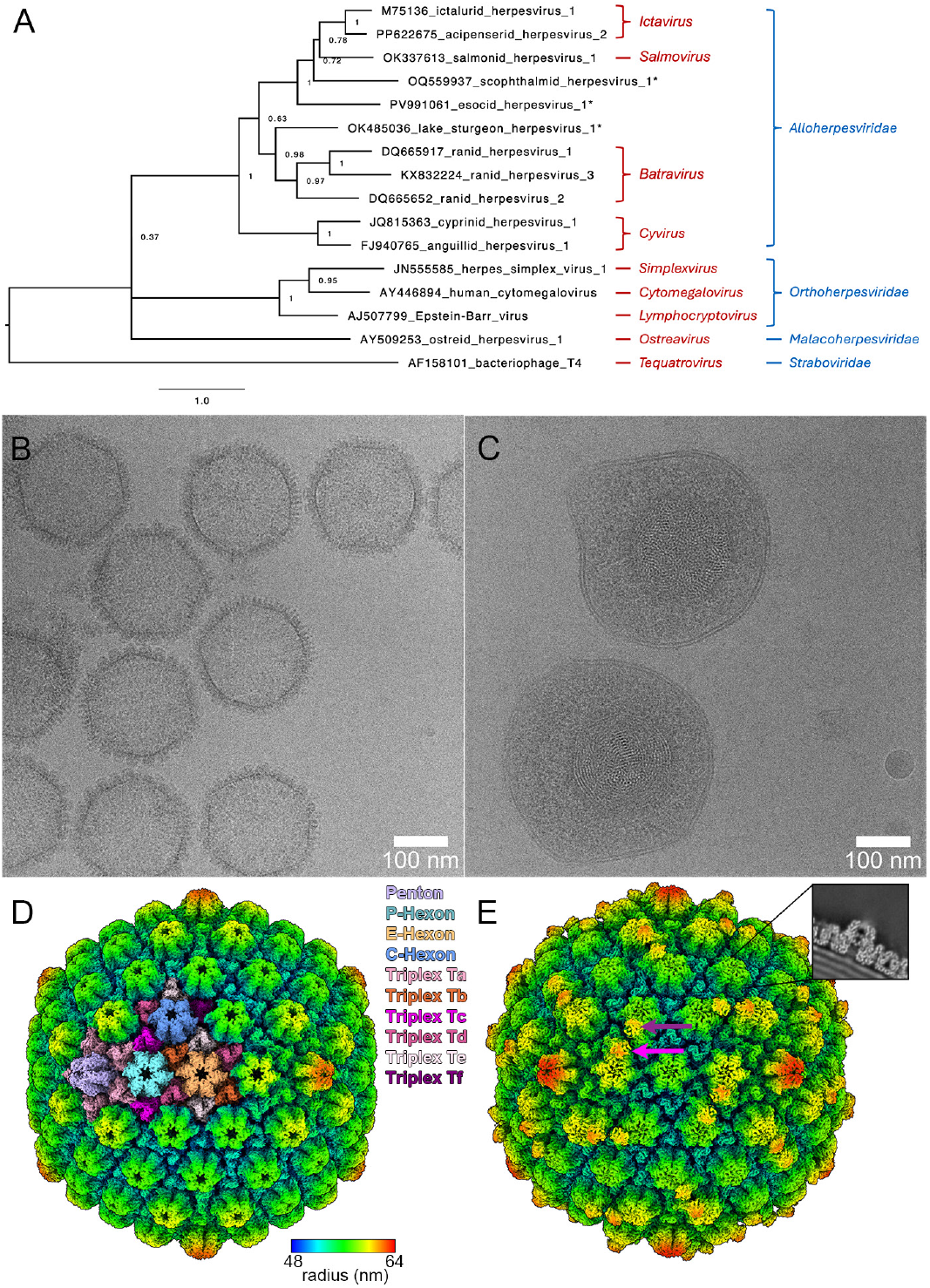
Phylogeny and Cryo-EM of IcHV-1. (A) Terminase based phylogeny of the *Alloherpesviridae* and related viruses. Amino acid sequence data for selected viruses representing the phylogenetic spread of herpesviruses were used along with that of the bacteriophage T4. The names of virus genera and families are in red and blue font, respectively. The nomenclature in black font consists of the underlying GenBank nucleotide accession number followed by the virus name, which is either a systematic one for classified members of the *Alloherpesviridae*, or a common one for unclassified viruses grouping in the *Alloherpesviridae*, and viruses in other families. The names of unclassified viruses are marked with asterisks. The scale shows the number of substitutions per position. (B,C) Micrographs of purified IcHV-1 capsids (B) and virions (C), scale bar = 100nm. (D,E) Icosahedral reconstructions of IcHV-1 capsid (D) and virion (E). Radial-depth cued isosurface representations are shown, viewed along a two-fold symmetry axis. The colour key indicates the radial distance in nm from the particle centre. The penton, hexons, and triplexes in the asymmetric unit are colour-coded and indicated in (D). Two copies of the inner-tegument protein pORF65 are indicated by purple/pink arrows in (E). In the virion reconstruction, density was found in the distal hexon channels, occluding the opening. Cross-sections through the reconstruction showed weak, blurred density radiating from this feature (inset, E).

Herpesviruses encode large (∼200nm), enveloped virions. They package their large double-stranded DNA genomes (up to 295 kbp) in a T=16 icosahedral capsid (*6, 7*). This is surrounded by a proteinaceous layer termed the tegument, which contains ∼20-40 different proteins that shape the cellular environment supporting initiation of infection. The capsid is made up of 11 pentameric and 150 hexameric capsomeres (pentons and hexons respectively) assembled from the major capsid protein (*8, 9*). Lying between the capsomeres at sites of local three-fold symmetry are triplexes, which are heterotrimers made up of one copy of the triplex-1 protein and two copies of the triplex-2 protein, and are thought to direct capsid assembly. Pentons are located at 11 of the 12 five-fold vertices in the icosahedral lattice, while at the 12^th^ - the portal vertex – there is a dodecameric portal assembly through which the viral genome is delivered into the capsid by an ATP-dependent terminase complex (*10-12*). Upon genome packaging, inner tegument proteins attach to the outer surface of the capsid, presumably to secure the packaged genome and strengthen the capsid as well as acting as a platform for the later addition of the remainder of the tegument. The pattern of tegumentation varies between different herpesviruses. In HSV, the capsid vertex specific components (CVSC – also known as capsid associated tegument complex CATC) and portal-vertex associated tegument (PVAT) complex, comprising pUL17, pUL25 and pUL36, assemble about the penton- and portal-vertices respectively, forming a star-shaped density. At the portal-vertex 10 copies of the C-terminal domain of pUL25 form a cap, comprising two pentameric tiers, that appears to bind the end of the packaged DNA lying in the portal channel (*10, 11*). The icosahedral tegument is more extensive in HCMV, forming a network over the entire capsid surface, perhaps providing additional reinforcement to support packaging of a larger genome (*9, 13, 14*). Capsid assembly, genome packaging and, in the case of the *Alphaherpesvirinae* HSV and VZV, addition of the CVSC, takes place in the host-cell nucleus (*15-17*). Mature nucleocapsids then bud across the nuclear membrane (*18*) and enter the cytosol where the remainder of the tegument is added prior to budding into cellular compartments derived from the trans-golgi network or plasma membrane, from which they acquire their viral envelope and associated viral glycoproteins. Finally, mature enveloped virions are secreted from the infected cell by exocytosis (*19*).

Ictalurid herpesvirus 1 (IcHV-1; species - *Ictavirus ictaluridallo1*) also known as channel catfish virus, is a member of the *Alloherpesviridae* and infects the channel catfish *(Ictalurus punctatus*) causing substantial economic impact to aquaculture industries. IcHV-1 was first isolated in the late 1960s from channel catfish exhibiting haemorrhagic disease associated with high mortality rates in fish-farms in the U.S. states of Alabama, Arkansas, Kentucky and Texas (*20*). The virus was subsequently shown to have herpesvirus-like morphology when viewed in the transmission electron microscope (*21*). However, analysis of the IcHV-1 136 kbp genome sequence did not reveal detectable conservation of herpesvirus protein sequences beyond the presence of enzymatic motifs also present in non-viral proteins and a T4 gp17-like terminase, similar to HSV pUL15 and its orthologues in other members of the *Orthoherpesviridae* (*22*). Mass-spectrometry analysis of purified IcHV-1 capsids and virions identified 11 structural proteins thus enabling assignment of the protein products of open reading frame 39 (pORF39) as the major capsid protein and pORF53 and pORF27 as the triplex 1 and 2 proteins respectively, pORF65 and pORF11 as inner-tegument proteins and pORF59 as the major envelope glycoprotein (*23*). A low-resolution cryo-EM reconstruction of purified IcHV-1 capsids showed that they have the T=16 pentamer/ hexamer clustered icosahedral capsid characteristic of all herpesviruses analysed to date. However the technological limitations of the time prevented further insights (*24*).

Here we present the cryo-EM structures for IcHV-1 empty capsids purified from infected cell nuclei and for DNA containing capsids within purified virions (hereafter referred to as capsids and virions respectively). We have used icosahedral reconstruction and particle-subtraction approaches to calculate maps for the symmetrically ordered penton and hexon capsomeres and triplexes at ∼3Å resolution in both the capsid and virion. Using symmetry breaking methods, we have calculated reconstructions for the capsid and virion portal vertices. These revealed a substantial PVAT complex in the virion with features that are quite different from those of human herpesviruses. We confirmed the identities and constructed atomic models for the capsid proteins pORF 39, pORF53 and pORF27 in both empty and full capsids. Further understanding at this point was limited by the general lack of detectable protein sequence homology between IcHV-1 and better characterised members of the *Orthoherpesviridae*, which led to the limited annotation of the IcHV-1 genome (Genbank accession number M75136.2) mentioned above. However, recently developed machine-learning software for *de novo* model building enabled the identification and modelling of the portal protein (pORF37), an icosahedrally ordered inner tegument protein (pORF65) that is also incorporated in the PVAT complex, a largely disordered inner tegument protein that occludes the hexon channel (pORF42) and a further four novel PVAT proteins (pORF23, 24, 66 and 67). Although clear conservation of structure is evident in the core capsid proteins (pORF39, 27 and 53), the dodecameric portal was found to have a rather different overall shape while maintaining structural motifs common to the known portals of both *Herpesvirales* and *Caudoviricetes*. Of the remaining six identified proteins, we found that pORF23 is an orthologue of the *Orthoherpesviridae* CVSC/PVAT protein pUL17, while pORF24, 42, 65 and 66 showed no compelling similarity to known protein structures. The most intriguing outcome of our analysis of these novel structural proteins was the finding that pORF67 comprises four domains that have macrodomain-like topology.

## Results

### Cryo-EM of IcHV-1 capsids and virions reveals conserved major capsid protein and triplex structures and two novel inner tegument proteins

To investigate the structure of IcHV-1, we purified both capsids and virions from infected brown bullhead cells. Capsids were isolated from infected cell nuclei and virions were purified from the culture medium. Both were prepared for cryoEM by plunge freezing and then imaged in a JEOL Cryo-ARM 300 equipped with a Direct Electron Apollo detector, at the Scottish Centre for Macromolecular Imaging (Figure 1B-C). Three-dimensional image reconstruction with imposition of icosahedral symmetry yielded intermediate resolution (∼5Å) maps of the symmetric features (Figure 1D-E, Movie S1). The capsid map confirms the previously reported T=16 icosahedral symmetry (*24*) comprising pentons at each five-fold vertex (owing to the imposition of icosahedral symmetry), peripentonal (P), central (C), and edge (E) hexons, and six unique triplex positions (Ta-Tf). As well as the presence of concentric shells of packaged DNA within the virion reconstruction there were two notable differences between the virion and capsid maps. In each asymmetric unit, two copies of a large globular tegument protein were clearly resolved (120 all together)., one on top of triplex Tb and leaning against the C hexon, and the other sat on top of triplex Tc and leaning against the E hexon. Furthermore, density was seen to occupy the distal sections of each hexon channel, extending out of the hexon tips as fuzzy weak density (Figure 1E inset).

Achieving only intermediate resolution maps for large icosahedral capsids is often a consequence of deviations from strict symmetrical equivalence that lead to blurring of high-resolution information by incoherent averaging. To overcome this limitation, we used particle subtraction and sub-particle refinement to reconstruct individual capsomeres in the asymmetric unit, thus achieving maps at resolutions of 3-3.1Å. To confirm the assignment of IcHV-1 pORFs 39, 53 and 27 as structural proteins and identify the novel inner tegument proteins, the sub-particle reconstructions were subjected to *de novo* model building using ModelAngelo without providing any sequence information (*25*). This approach yielded atomic models based solely on the cryo-EM density, comprising extensive regions of contiguous sequence. Chains were selected within the hexon, the triplex and the larger inner tegument protein and submitted to the protein BLAST server (*26*) (https://blast.ncbi.nlm.nih.gov/Blast.cgi). In all cases the topmost hit was for an IcHV-1 protein with sequence identity values ranging between 58% and 75%, and with 100% coverage. Unbiased searching of the NCBI protein sequence database, rather than searching only predicted proteins annotated for the IcHV-1 genome, lends confidence to our assignment of identities to each component of the capsid and virion. These data confirmed that the major capsid protein is encoded by ORF39, triplex 1 by ORF53 and triplex 2 by ORF27. Furthermore, we identified the large globular inner tegument protein as being encoded by ORF65. The density located in the hexon channel was too small to identify using the blastp approach but was identified as pORF42 using the built in HMMER (*27, 28*) algorithm in ModelAngelo to search against the full IcHV-1 proteome. The predicted sequences of the proteins identified were then used to build models for the asymmetric units of both the capsid and the virion (Figure 2A-C)).

**Figure 2.**
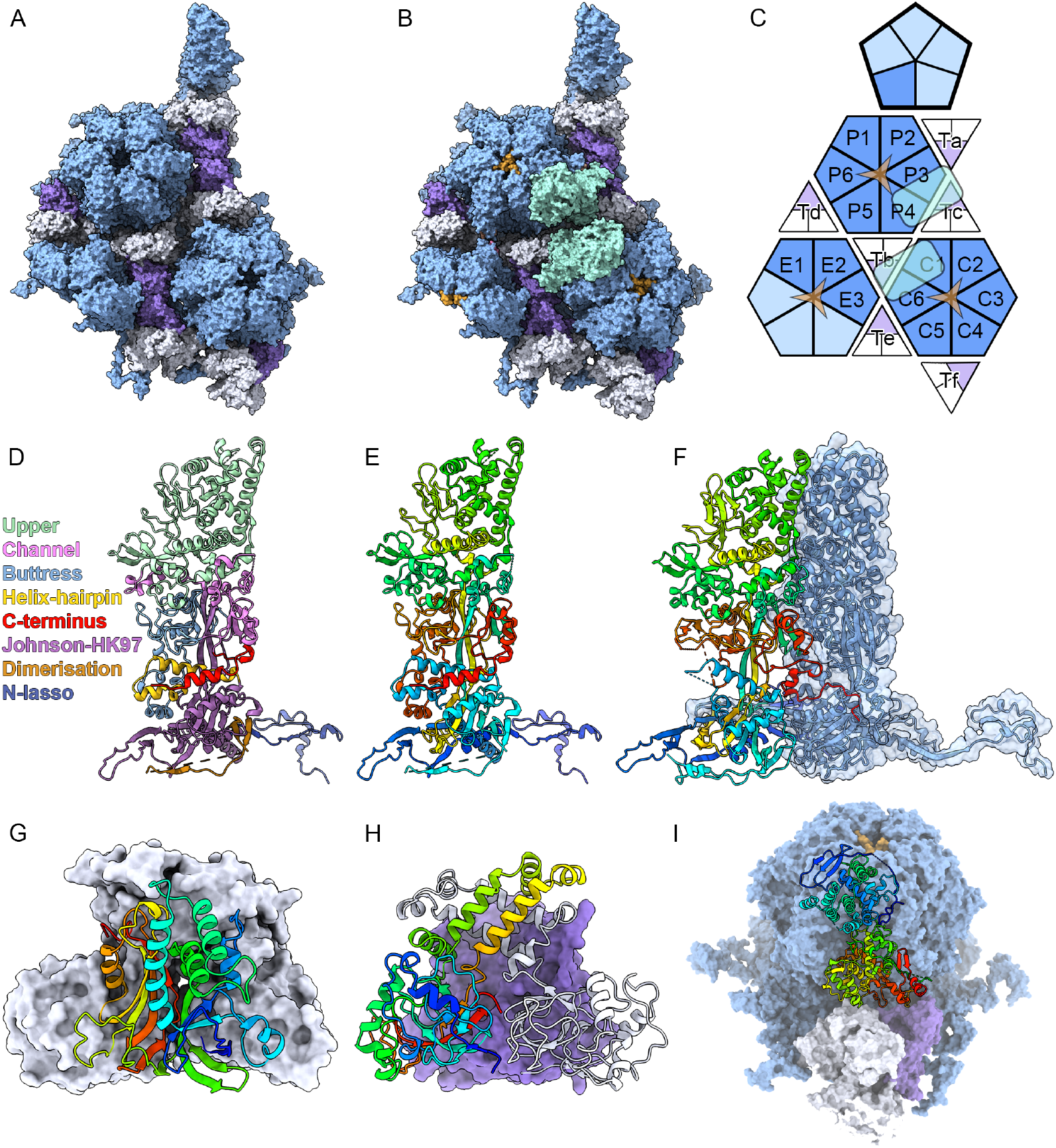
Sub-particle reconstruction enabled high-resolution structure determination of the icosahedral asymmetric units of both capsid and virion. Solvent-excluded surface (SES) representations of the asymmetric units of the IcHV-1 capsid (A) and virion (B). The asymmetric units contain 16 copies of the major capsid protein pORF39, (blue): six (P1-P6) make up a complete P-hexon, six (C1-C6) make up a complete C-hexon, and three (E1-E3) contribute to half of an E-hexon. One copy is present and contributed to the penton. The asymmetric unit comprises six triplexes each composed of one copy of pORF53 (purple), and two copies of pORF27 (white). In the virion, two copies of the inner-tegument protein pORF65 are positioned on the Tb and Tc triplexes. Additionally, three copies of pORF42 occlude the distal tips of the hexon-channels. (C) Schematic representation of the virion asymmetric unit coloured as in (A)–(B). (D) A ribbon diagram of the penton pORF39 is shown, color-coded to highlight the domain structure previously defined for orthoherpesvirus MCPs. The IcHV-1 MCP features an extended C-terminal region (red), which folds toward the helix–hairpin domain to form a helical bundle. This region and adopts different conformations in penton versus hexon environments (compare E and F). (E) A ribbon diagram of the penton MCP is shown in rainbow colour-scheme from N-terminus (blue) to C-terminus (red). (F) A ribbon diagram of the hexon MCP is shown in the same rainbow colour-scheme. A second, neighbouring copy is shown in both ribbon and transparent sovent-excluded surface (blue), to highlight the domain insertion of the C-terminal domain in this form of the IcHV-1 capsomere. (G) A ribbon diagram of pORF53 is shown in rainbow colour-scheme, with the associated two copies of pORF27 shown as SES (white). (H) Two copies of pORF27 are shown as ribbon diagrams, one in rainbow colour-scheme and one in white. The associated copy of pORF53 is displayed as a purple SES. (I) SES representation of a P-hexon (blue/gold) and adjacent Tc triplex (purple/white). pORF65 is shown as a ribbon diagram in rainbow colour-scheme.

### Structure of the major capsid protein pORF39

The structure of the 123kDa major capsid protein pORF39 shows a high degree of conservation when compared with the major capsid proteins of other known herpesviruses (Figure 2D-F, Movie S1). Previous studies have defined seven domains for herpesvirus major capsid proteins (*9*). In IcHV-1 pORF39 these are the upper domain (483–833), channel domain (394–482 and 1053–1086), buttress domain (910–1052), helix-hairpin domain (205–242), Johnson/HK97 fold (85–204, 243–281, 355–393, and 834–910), dimerisation domain (282–354), and N-lasso (1–84) (*9*). These domains form three sections: the upper section, middle section (channel, buttress, and helix-hairpin), and lower section (Johnson-fold, dimerisation, and N-lasso). The upper and middle sections form the turret-shaped capsid protrusions of the hexon and penton capsomeres, whereas the lower section forms the capsid floor in which pORF39 protomers are laced together through extensive interactions that closely follow those described previously in orthoherpesviruses (*8, 9*). Intracapsomere interactions between neighbouring protomers are stabilised by β-sheet augmentation between the N-lasso and Johnson/HK97 domains, and longer-range N-lasso and dimerisation domain interactions extend between capsomeres.

Structural comparison of hexon and penton forms revealed similar conformations in the upper section, moderate alterations in the middle section, and substantial conformational differences in the lower section. In the penton, the Johnson-fold and N-lasso undergo an approximately 15° counterclockwise rotation, which straightens the subunit and increases the curvature of the penton floor. The helix-hairpin domain and buttress domain, also undergo a reorganization, transitioning from a more flexible conformation in the hexon, identified by regions of poorly resolved density in the cryoEM map, to a well-ordered state in the penton. Interestingly, the C-terminus of pORF39 from amino-acid residue (aa) 1087 undergoes a substantial reorganisation between penton and hexon forms and for this reason we have identified it as a distinct domain. In the penton the last resolved aa is 1111, and aa 1101-1111 form an α-helix that is oriented approximately parallel to the capsid floor to form a four-helix bundle with the helix-hairpin and HK97 domains. In the hexon, the C-terminal aa from 1087 turn in the opposite direction (clockwise as viewed from the capsid exterior), and insert into the neighbouring hexon subunit, occupying the same position above the helix-hairpin motif. This C-terminal domain-swap has not been observed in structures of orthoherpesvirus major capsid proteins and may provide an additional stabilising interaction between pORF39 subunits in IcHV-1 hexon capsomeres or play a role in regulating capsid assembly.

### Structure of the triplex proteins pORF53 and pORF27

Like the major capsid protein, IcHV-1 triplex proteins show an overall conserved architecture when compared to that in the. The heterotrimeric complex comprises one copy of pORF53 (triplex 1) and two copies of pORF27 (triplex 2) (Figure 2G-H). Occupying sites of local three-fold symmetry between the capsomeres, triplexes likely stabilise the capsid floor, bracing the capsid against the considerable internal pressure from the packaged genome. The pORF27 dimers have additional stabilising interactions through their ‘embracing arms’ at the distal tips of the triplex. Our cryo-EM maps supported the modelling of the entirety of pORF27 – aa 1-288, but the C-terminal 23 aa are not resolved for pORF53. Peri-pentonal Ta triplexes are oriented such that both copies of pORF27 are oriented towards the penton, whereas at the portal vertex the Ta triplexes are rotated such that one copy of pORF53 and one copy of pORF27 are oriented to face the five-fold symmetry axis (see below).

### The inner tegument proteins pORF65 and pORF42 are partially unresolved

A novel feature of the IcHV-1 virion structure is the presence of two copies of the inner tegument protein pORF65, in each asymmetric unit. This 1434 aa protein is the second largest encoded by IcHV-1 (after pORF67), but neither copy is completely resolved in our cryo-EM map. Indeed, only aa 14-720 are modelled in the copy associated with the Tb-triplex and C-hexon, and aa 17-720 are modelled in the copy associated with triplex Tc and the P-hexon. In both models there are multiple short chain breaks. Thus, the structure for a substantial proportion of this protein is not determined. The resolved structure presents a globular protein comprising two domains, with the C-terminal domain comprising aa 352-720 proximal to the capsid surface and resting on the underlying triplex, and the distal N-terminal domain (aa 14/17-351) leaning between two pORF39 upper domains at positions P3 and P4 in the P-hexon and between protomers C1 and C6 in the C-hexon (Figure 2C).

Structural comparison of IcHV-1 hexons with those from orthoherpesviruses revealed a markedly narrower axial channel. In IcHV-1 virions the hexon channel vestibule is sealed by three copies of a short helical density. In our cryo-EM maps the density suggests six copies with partial occupancy, however modelling this feature shows that a maximum of three copies of pORF42 can occupy the available space at alternating sites owing to steric collision. This suggests that the arrangement of pORF42 within the six identified binding sites inside a channel is not necessarily determined by icosahedral symmetry and each hexon within a virion may have one of several possible arrangements. The blurred density extending from the hexon channel in the map suggested that the hexon channel density is likely part of a larger but poorly ordered inner tegument protein. Owing to the small portion of density observed, identifying this peptide was only possible using the ModelAngelo HMMER tool as noted above. Following the combined averaging of the channel density from the three hexons in the asymmetric unit, density for several bulky sidechains was resolved sufficiently well as to lend confidence to our assignment of this feature as aa 2-17 of pORF42 (Movie S1). Further supporting this assignment, a predicted structure of pORF42 calculated using the machine-learning tool colabfold (*29*) within the Viro3D database (https://viro3d.cvr.gla.ac.uk) shows a largely disordered protein, with the only secondary structure being an N-terminal α-helix (aa 4-23) (*30*). The hexon channel density was only observed in the mature virion and may serve as a platform for subsequent tegument addition, perhaps with the disordered region supporting interactions with multiple binding partners. Hexons from orthoherpesviruses typically exhibit a more open channel and sometimes are decorated with additional capsid-associated proteins at their distal tips (VP26 in HSV, SCP in VZV, and pBFRF3 in EBV) (*8, 31, 32*); these accessory components are absent in IcHV-1.

### Focussed classification reveals a large and elaborate ‘tail’ at the virion portal-vertex

To resolve the portal vertex for the capsid, we applied symmetry expansion and focussed classification (*11*). Identification of the location of the portal vertex in each capsid particle image facilitated calculation of a C5 symmetric reconstruction of the capsid at 7Å resolution in which the portal vertex was aligned to the z-axis of the map (Figure 3A-C). The capsid portal was resolved as noisy density that was not tightly apposed to the opening in the portal vertex. Poorly resolved portal density is a consequence of a symmetry mismatch between the portal and capsid; all herpesvirus portals studied to date have been shown to have C12 symmetry. Furthermore, in the capsid, the portal does not appear to be firmly constrained by interactions with the capsid shell. Aside from the portal, no other features were observed at the capsid portal vertex.

**Figure 3.**
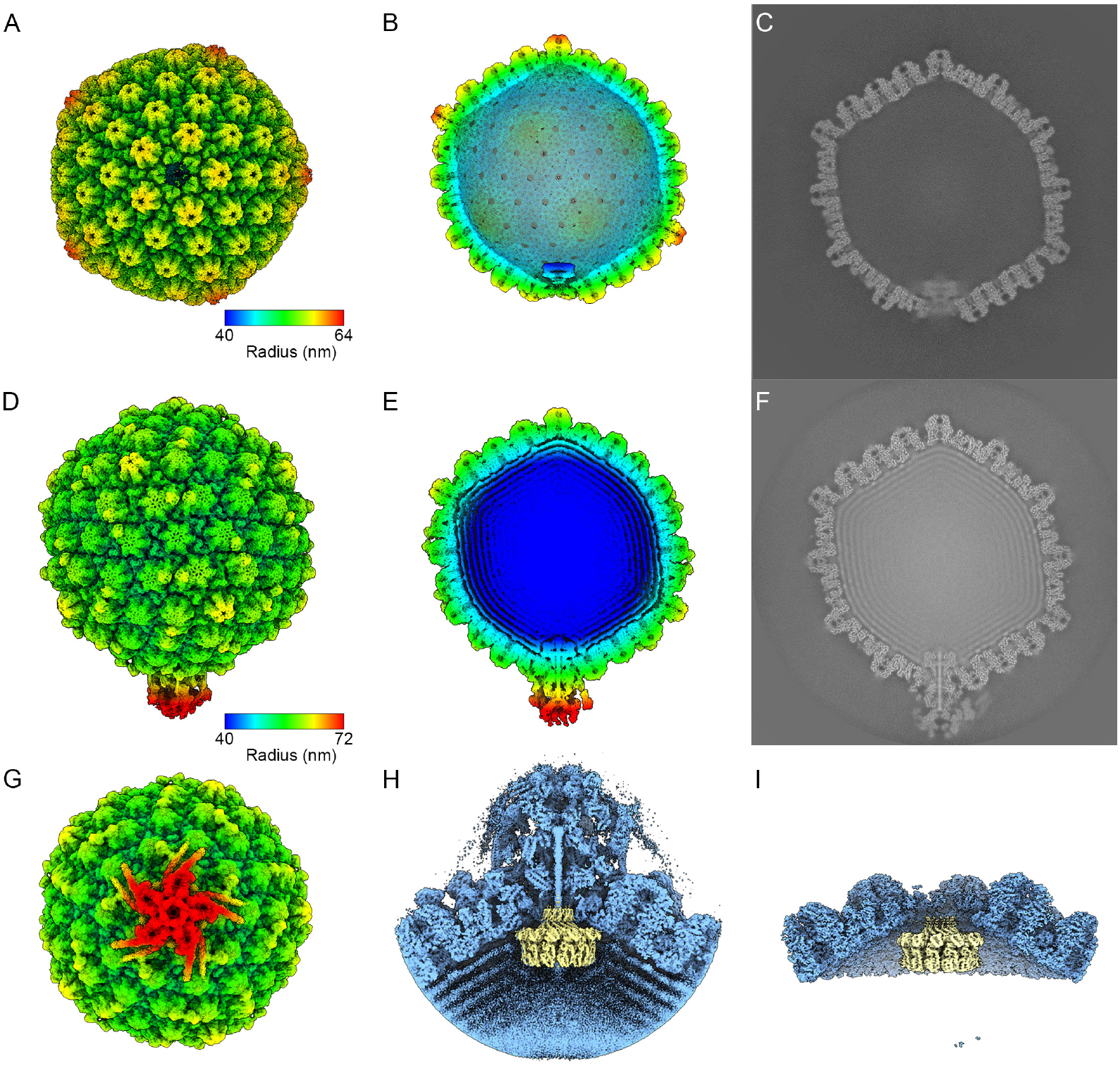
Symmetry breaking reveals the structure of the capsid and virion portal vertex. (A) Three-dimensional reconstruction of the C5-symmetric capsid following symmetry expansion and focussed classification to reveal the portal vertex. The map is shown as a radial-depth cued isosurface view along the portal-vertex, coloured according to distance from the particle centre (colour key). (B) Cut-away view of the C5 symmetric capsid reconstruction viewed perpendicular to the C5 symmetry axis of the portal-vertex, revealing the capsid interior and the symmetry-mismatched portal density at the interior of the portal-vertex (bottom-centre). (C) A central slice through the C5 capsid reconstruction showing the blurred, symmetry mismatched portal density at the portal-vertex interior (bottom-centre). (D) Three-dimensional reconstruction of the C5-symmetric virion following symmetry expansion and focussed classification to reveal the portal vertex. The map is shown as a radial-depth cued isosurface view perpendicular to the C5 symmetry axis of the portal-vertex, coloured according to distance from the particle centre (colour key). The large portal-vertex associated tegument complex is visible at the bottom of the particle. (E) Cut-away view of the C5 symmetric virion reconstruction viewed in the same orientation as (D), showing the packaged DNA arranged in dense, concentric shells. The PVAT and portal density are clearly seen as a large tail-like complex at the bottom of the particle. (F) A central slice through the C5 virion reconstruction showing the symmetry mismatched portal density at the portal-vertex interior and the large tail-like PVAT complex (bottom/centre). (G) View of the C5-symmetric virion map viewed along the portal-vertex axis at a reduced isosurface threshold, showing the weak, noisy radial fibres extending from the PVAT. (H) Cut-away view of the sub-particle C5 reconstruction of the virion portal-vertex (blue) with the C12 reconstruction of the portal superimposed (yellow). (I) Cut-away view of the sub-particle C5 reconstruction of the capsid portal-vertex (blue) with the C12 reconstruction of the portal superimposed (yellow).

Taking the same approach, we calculated a C5 symmetric reconstruction of the virion at 5Å resolution. This revealed the presence of an unexpected and substantial PVAT assembly at the portal vertex exterior (Figure 3D-H). The large, complex PVAT density is superficially reminiscent of the short non-contractile tails seen in phages of the *Podoviridae* (*33*), extending some 28 nm from the floor of the capsid shell. The map shows that the portal complex is pressed into the portal vertex aperture, presumably by the pressure of the packaged genome. The features of the portal are better resolved owing to it being constrained by close contact with the capsid shell, although the symmetry mismatch still prevented assembly of an atomic model. Running through the centre of the portal and PVAT complex is a rod of density 27nm in length. In human herpesviruses this density has been postulated to be the trailing end of the packaged genome, an assignment that would seem likely here although we cannot exclude the possibility that this feature is protein. The PVAT density appears well ordered in the regions closest to the portal, being sharply resolved and revealing the presence of extensive coiled-coil secondary structure. PVAT features become less sharp in the distal section of the map presumably due to flexibility in the assembly. When viewed in section (Figure 3F) blurred density radiating from the PVAT is evident. These weaker features can be visualised in 3D isosurface representation by setting a low threshold value (Figure 3G). Three unique radial fibres extend 22nm suspended over the capsid surface (15 altogether considering the C5 symmetry). Two fibres are stacked one on top of the other with the inner-most one having a larger diameter and terminating above the P hexon while the outer-most one extends towards the E hexon. The third fibre is displaced counterclockwise at an angle of 14 degrees and rests on top of the P-hexon-bound copy of pORF65, also extending towards the E-hexon.

### ModelAngelo enables identification and modelling of the portal protein at resolution poorer than 4 angstroms

To achieve sufficient resolution to support the construction of an atomic model of the portal and PVAT, particle subtraction and subsequent local refinement of the portal vertices was performed for both the capsid and virion. This yielded maps with improved global resolution estimates of 4.4Å for the capsid and 4.1Å for the virion. To resolve the symmetry mismatch between the portal vertex (C5) and the dodecameric portal, we performed focussed classification with local refinement and imposition of C12 symmetry applying a mask that enclosed only the portal density. This led to the calculation of portal maps for both the capsid at 4.3Å resolution, and virion at 4.6Å resolution (Movie S2, Figure 3H-I). Despite the modest resolution achieved, ModelAngelo was able to build a high-quality model in the absence of sequence data, using the capsid portal reconstruction. Submitting sequences from the built model for protein blast analysis identified the portal protein as pORF37. Interestingly pORF37 was identified as having a similar fold to the portal proteins of alpha-herpesviruses (pUL6), by machine-learning based structure prediction (*29*) combined with foldseek analysis (*34*) within the Viro3D database (*30*).

Providing the sequence of pORF37 to ModelAngelo and subsequent manual editing and refinement led to high-quality models for both capsid and virion portals (Figure 4A-B). The IcHV-1 portal exhibits an overall architecture similar to that of known *Orthoherpesviridae* and *Caudoviricetes*. Herpesvirus portal proteins are described as having six domains. Moving from the virion exterior inwards, these are the turret, clip, stem, wing, β-hairpin/tunnel-loop, and wall. In the deposited coordinates for the C12 symmetric elements of pUL6 (PDB 6OD7 (*10*) – Figure 4C) – the portal protein of HSV-1, the turret is not resolved (aa 308-516) however the C5 virion reconstruction shows that it includes ‘tentacle-helices’ that form a truncated-conical assembly that replaces the penton capsomere at the portal vertex (*10, 11*). The turret is inserted between the clip (aa 301-307), which traverses the aperture in the capsid shell, and the stem which has two α-helices (aa 272-300 and 517-540) and connects the clip/turret to the wing and β-hairpin domains. The wing (aa 33-62 and 150-271) braces against the inner surface of the capsid, while the β-hairpin (aa 541-558) engages the DNA running through the centre of the portal DNA translocation tunnel. The remainder of the inner surface of the portal DNA translocation tunnel comprises the wall domain (aa 63-149 and 559-623). Several notable differences are apparent when comparing the IcHV-1 portal to that of HSV. In IcHV-1 the portal presents a wheel-shaped density, whereas orthoherpesvirus portals resemble a bowl with its opening oriented towards the capsid interior. The pORF37 turret (aa 366-505) also resolves the symmetry mismatch between the C12 portal and C5 portal-vertex through the formation of tentacle helices, but these are splayed out to engage with the peri-portal Ta triplexes (see below). The dodecameric portal structure follows a broadly similar clip, stem, tunnel-loop, wing and wall structure. The clip domain (aa 355-365 and 506-529) features a three-stranded β-sheet, with one β-strand donated from a neighbouring pORF37 protomer (Figure 4I). The ‘tunnel-loop’, which features a β-hairpin and engages the terminal DNA that extends through the portal-channel in several herpesvirus and phage portals, is not fully resolved in our structure and there is no clearly resolved contact between the portal and the DNA rod laying at the centre of the C12 map. Presumably, the tunnel loop (550-572) is not stabilised by an interaction with the end of the packaged genomic DNA. The stem domain (aa 328-354 and 530-549) bridges the clip to the wing and tunnel-loop. As the pORF37 wing (aa 45-327) and wall (aa 573-643) domains are folded rather differently than in orthoherpesvirus portal proteins, forming a more compact assembly, we have defined them as the N-terminal and C-terminal regions respectively.

**Figure 4.**
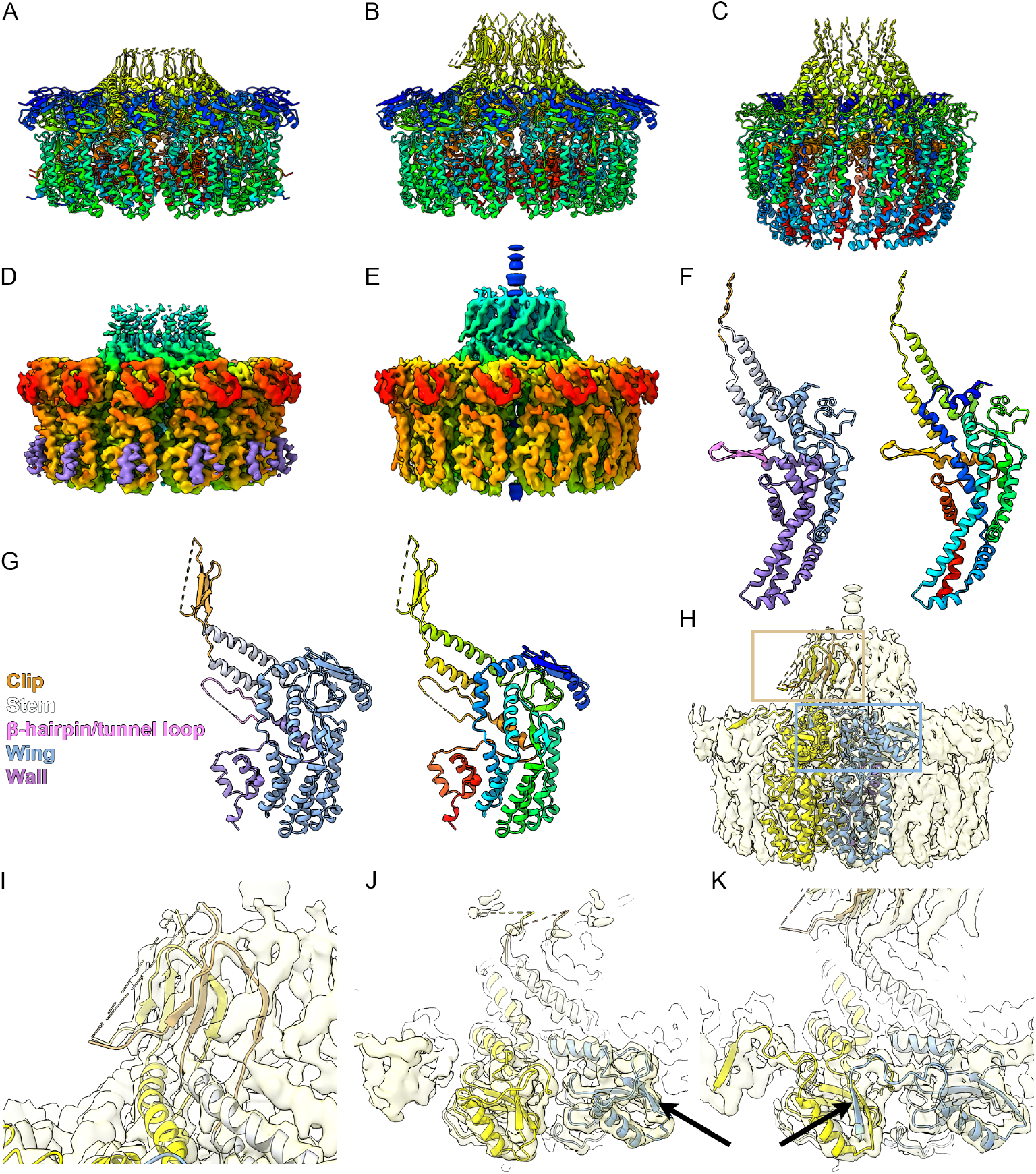
Architecture of the capsid and virion portal. (A–C) Ribbon diagram representations of the portal complexes of the IcHV-1 capsid (A), virion (B), and that of HSV-1 (C), each shown in rainbow colouring. The IcHV-1 portal consists of 12 copies of pORF37, likewise the HSV-1 portal complex comprises 12 copies of pUL6. All structures exhibit C12 symmetry. (D, E) Corresponding cryo-EM density maps of the IcHV-1 capsid (D) and virion (E), portals, resolved to 4.3 Å and 4.6 Å respectively. In the capsid map, an additional density is observed on the outer face of the wing domain (purple), assigned by ModelAngelo as the N-terminus of pORF37. Due to discontinuity in the density, we were unable to determine whether this short α-helix originates from the same monomer or is contributed by a neighbouring subunit. (F, G) Structural comparison of HSV-1 pUL6 (F) and IcHV-1 Orf37 (G). Each monomer is shown as a ribbon diagram coloured to highlight the domain structure (left) and rainbow representation (right). (H) Two adjacent virion portal pORF37 monomers are shown as ribbon diagrams (yellow and blue) within the reconstructed volume (transparent surface), to highlight two instances of β-augmentation. Yellow and blue boxes indicate unique monomer–monomer interaction regions. (I) Enlarged view of the yellow-boxed region, showing β-strand augmentation in which a three-stranded β-sheet is assembled by two neighbouring protomers to form the clip region. (J, K) View of the top of the wing domain in the capsid (J) and virion (K) portal, highlighting a rearrangement between the two forms whereby the N-terminal β-strand which contributes to a four-stranded β-sheet in one monomer (black arrows) inserts instead into the same β-sheet in the neighbouring protomer, forming an inter-subunit β-sheet extension. This rearrangement likely accounts for the loss of the N-terminal helical density at the base of the wing domain in the virion portal.

Comparison of the capsid and virion portal maps shows that pORF37 undergoes a substantial rearrangement in the wing domain upon DNA packaging. In the capsid form of the portal, aa 45-62 fold over the top-most part of the wing contributing a β-strand to a six-stranded β-sheet (Figure 4J). In the virion however, this region inserts into the neighbouring protomer donating the β-strand to form the same six-stranded β-sheet in that molecule (Figure 4K). A second difference in the N-terminal region is observed towards the bottom of the wing domain in the capsid portal, where a small section of helical density is present (coloured purple in Figure 4D) that was not resolved in the virion portal. The predicted structure for pORF37 in the Viro3D database, together with the rearrangements we observed between the capsid and virion forms of pORF37 to accommodate the β-augmentation in the wing domain, led us to assign this density to aa 4-18 of pORF37. A comparable N-terminal helix-loop motif has been described in the portal protein of HCMV pUL104 (*35*), where it is referred to as both an anchor and the aileron domain, owing to it being positioned over the wing domain. However, in that case the N-terminal element is located at the top of the wing domain, atop a circular fragment of DNA that has been reported in all portal structures from human herpesviruses studied to date. Our data do not show the presence of such a DNA component encircling the IcHV-1 portal.

### Structure and composition of the PVAT tail

Although the PVAT presented noisy, poorly resolved density in the distal portions of the C5 sub-particle reconstruction, we were able, through a process of masking and local refinement, to prepare a series of reconstructions that ranged in resolution between 4.1Å and 4.5Å. These supported the identification and modelling of a substantial proportion of the assembly. Using the same approach of supplying ModelAngelo with a map only and submitting the sequences from the resulting models to the BLAST server, we were able to identify four additional IcHV-1 proteins as components of the PVAT complex: pORF23, 24, 66 and 67 (Figure 5, Movie S2).

**Figure 5:**
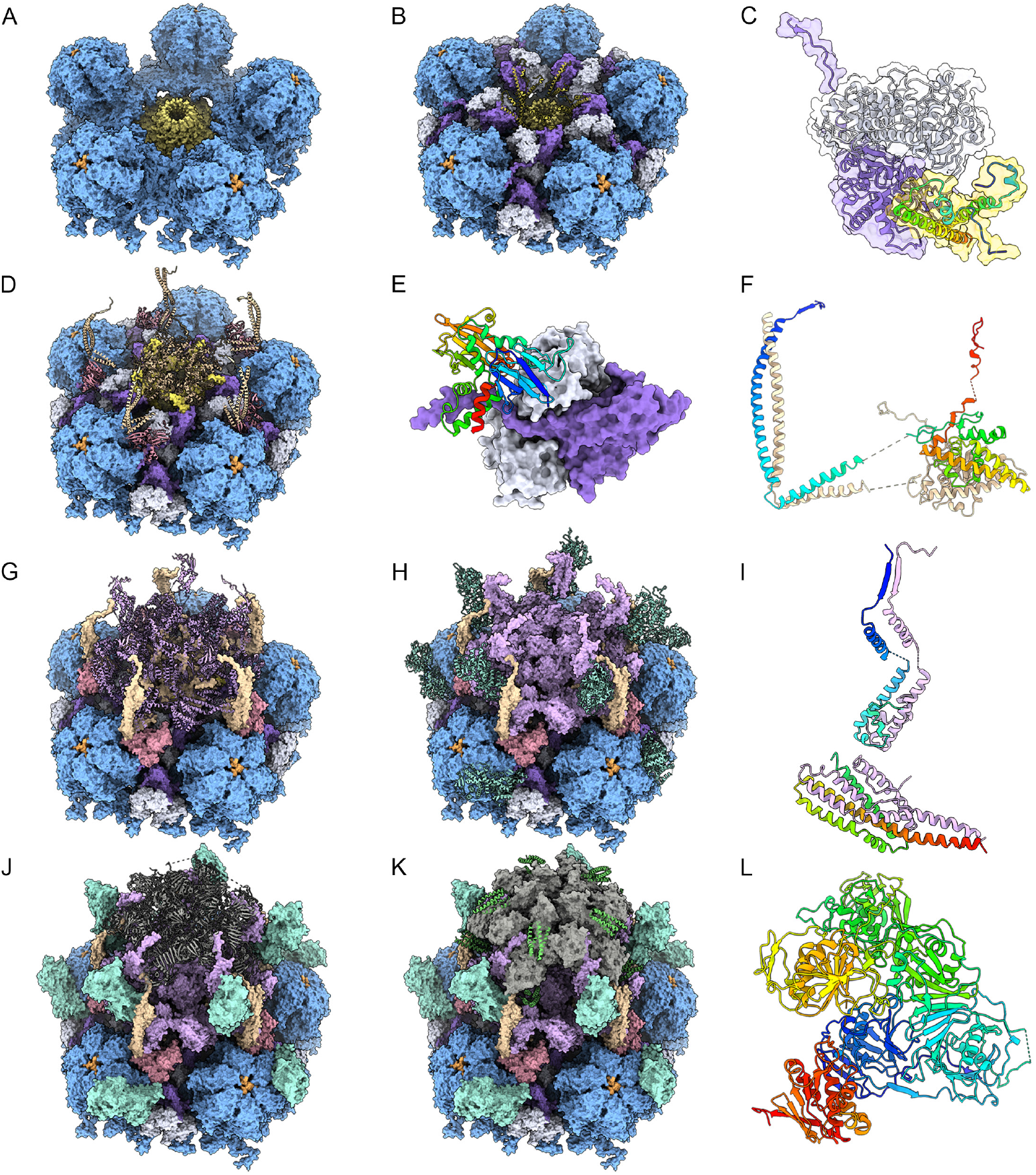
Architecture of the Portal Vertex Associated Teguments complex. (A–L) Each panel shows a stepwise addition of PVAT components, or views of individual proteins. Newly added proteins are displayed as ribbon-diagrams, while previously positioned components are shown as solvent-excluded surface (SES) representation. (A) Five P-hexons (SES, blue) arranged with C5 symmetry about the portal vertex. The underlying portal complex is shown (SES, yellow). pORF42 is shown occluding the vestibule of the hexon channels (SES, orange). (B) Five Triplexes (pORF53 in purple, pORF27 in white; SES) are oriented such that pORF53 presents a cleft to engage five turret tentacle-helices of pORF37 (ribbon, yellow) that extend from the portal. Five additional pORF37 turret tentacle-helices are adopt a second conformation such that ten copies resolve the C5/C12 symmetry mismatch. (C) A close-up view of the two conformations of the pORF37 tentacle-helices is shown (rainbow coloured ribbon diagram, SES transparent yellow), binding to pORF53 (purple ribbon and SES). pORF27 is shown to complete the Ta triplex (white ribbon and SES). (D) Ten copies each of pORF23 (ribbon, pink) and pORF24 (ribbon, wheat) form the next tier of the PVAT. The N-terminal ‘upstand’ of pORF24 rests on top of pORF23, while the C-terminal domain of pORF24 engages the pORF27 tentacle-helices to form a collar around the opening in the capsid floor at the portal-vertex. (E) pORF23 (ribbon, rainbow) is a two-domain protein that sits astride a cantilevered N-terminal arm of the Ta triplex pORF53 (purple, SES) and pORF27 (white, SES). (F) Two pORF24 monomers assemble as a dimer, with the N-terminus projecting to form an upstand perpendicular to the capsid floor (pORF24 copy1 rainbow, ORF24 copy 2 wheat). (G) Thirty copies of pORF66 (ribbon, lilac) come together, forming dimers and making up a considerable part of the PVAT complex assembled onto underlying pORF23 and pORF24. (H) In addition to the icosahedrally ordered pORF65 inner-tegument proteins, five copies of pORF65 (ribbon, cyan) contribute to the PVAT complex binding predominantly to the N-terminal helices of pORF24. (I) A pORF66 dimer is shown in ribbon representation one copy is shown in rainbow colour scheme, the other in lilac. (J) Five copies of pORF67 (ribbon, dark grey) form the distal PVAT. (K) Additional four-helix bundles (ribbon, green) were modelled but could not be confidently assigned, these likely anchor the poorly resolved radial fibres to the PVAT. (L) Ribbon diagram of a pORF67 monomer, shown in rainbow colour scheme.

The IcHV-1 portal turret comprises “tentacle-helices” that resolve the symmetry mismatch between the C12 portal and the C5 portal vertex. Ten α-helices from pORF37 are resolved extending from the capsid interior through the opening of the portal vertex in two conformations (Figure 5B-C). The first comprises pORF37 aa 388-461 and the second consists of aa 418-468. The peri-portal Ta triplexes are rotated compared to those at the penton vertices (where the pORF27 dimer is oriented towards the penton), such that pORF53 presents a cleft to engage conformer one. The second pORF37 tentacle helix conformer contacts pORF53 minimally. Both conformations engage pORF24 (Figure 5D,F), which forms a star-shaped assembly comprising five dimers arranged about the exterior of the opening in the capsid shell at the portal vertex. The N-terminal aa 1-105 extend radially from the globular C-terminal domains as α-helices running parallel to the capsid surface (74-105), before turning through ∼90º to form upstanding α-helical coiled coils (aa 1-73), oriented perpendicular to the capsid shell. The connecting density between the C- and N-terminal domains is not well resolved and therefore has not been modelled. The C-terminal domain of pORF24 (aa 114-289) forms the base of the central channel through which the DNA is transferred in to- and out of-the capsid. This region appears to constrict the DNA translocation channel and possibly prevents premature genome release.

The N-terminal domains of pORF24 rest upon another PVAT protein, pORF23, which in turn rests upon a bridge formed by a C-terminal arm of Ta Triplex 1 (pORF53). In all other triplexes resolved in our study, the C-terminal 23 aa of pORF53 are not detected. However, here we observe the C-terminal aa 285-308 forming a cantilevered arm that inserts into the upper domain of pORF39 (MCP) at position P3 of the peri-portal P-hexon. pORF23 sits astride this arm. Our data enabled us to model aa 81-377 of this 418 aa protein (Figure 5E).

Above the star-shaped platform formed by pORF24 and contained within the cage formed by its N-terminal upstands, 30 copies of pORF66 arranged as 15 dimers make up a considerable portion of the PVAT density (Figure 5G,I). Despite contributing substantially to the mass of the PVAT – a considerable portion of the 411 aa pORF66 is not resolved in our map and were not modelled. At the outermost base of this section of the PVAT, the C-termini of one pORF66 dimer contact pORF23 extending as a two-helix coiled coil (aa 368-402) over the parallel helices of pORF24 (aa 74-104), they then fold back on themselves to form a six-helix bundle (aa 273-367). One region (aa 242-272) is not resolved in our cryo-EM map, but aa 148-241 extend towards the distal PVAT, forming a third globular α-helical domain (aa 188-230) terminating in an α-helix (aa 161-170) and a short β-strand (aa 155-160). This β-strand contributes to a 5-stranded parallel β-sheet formed from one other pORF66 dimer and a single strand donated by pORF67 (aa 256-260). The second dimer that contributes two β-strands to this domain has a similar overall structure, although the two-helix and six-helix bundles are oriented roughly perpendicular to the capsid shell, resting on the C-terminal domain of pORF24. Furthermore, the region connecting the C-terminal and N-terminal sections (aa 242-272) is well resolved, showing formation of a three-stranded β-sheet. In this dimer, more of the N-terminal globular domain is resolved (aa 152-250) and appears to engage the central rod of density that we attribute to DNA. The final pORF66 dimer stands on top of the Ta triplex pORF53 and only the C-terminal section is resolved (aa 190-411), although unlike those in other pORF66 chains – this section is completely modelled. However, the resolution of the N-terminal domain (aa 190-250) was not sufficiently adequate to allow a model of this domain to be built based on the density alone. Instead, this domain was flexibly fit into density with secondary structure restraints based on the N-terminal domain of the inner-most pORF66 dimer, which was conformationally very closely matched.

In addition to the icosahedrally-ordered pORF65 chains that occupy every asymmetric unit of the virions, we also found five copies of pORF65 contributing to the PVAT complex. Our map showed a large globular density on the outer side of the PVAT assembly. Although this region was not resolved at sufficiently high resolution to identify and model *ab initio*, almost all secondary structure elements of pORF65 matched accurately to low resolution density counterparts when the 4.5Å map of the portal vertex was convolved with a gaussian filter of width 1.3Å. Given the presence of secondary structure elements consistent with pORF65, we incorporated this model into the PVAT by fitting as a rigid-body, with the N-terminal domain oriented towards the capsid surface. In our model this pORF65 chain primarily contacts the N-terminal upstands of pORF24, making only minor contacts with pORF66 (Figure 5H).

At the apex of the DNA translocation channel, we identified another unique IcHV-1 PVAT protein, the largest protein encoded by IcHV-1, pORF67 (Figure 5J,L). Here, five copies of this 1556 aa protein form a large cap on the PVAT assembly that is constricted at its base by a β-sheet loop motif at aa 870-895 marking the point in the cryo-EM map where the central rod of density that we attribute to the end of the viral genome terminates. The adjacent pORF66 domains (aa 152-250) may also play a role in sealing the PVAT and preventing genome release. Despite being at the distal tip of the PVAT, and being poorly resolved in initial sub-particle reconstructions, through careful masking and local refinement we were able to calculate a reconstruction at sufficient resolution to model most of this intriguing protein (aa 50-484, 498-1148, 1287-1468).

Bound to pORF67, we observed two additional densities corresponding to bundles of four α-helices (Figure 5K – coloured green). At the available resolution, it was not possible to discern an identity for these PVAT components. The locations of these helical bundles suggest that they are the attachment points of the poorly resolved radial fibres shown in figure 3G.

### pORF23 is an ortholog of pUL17

To determine whether the newly identified PVAT components pORF23, 24, 65, 66, and 67 present novel folds, we used the DALI server to search for similar structures in the protein data bank (*36*). To evaluate the results, we considered DALI’s Z-score, and the RMSD of the aligned folds. We also evaluated the fold similarity qualitatively by visual inspection in ChimeraX. Where proteins were found to share similar folds with previously solved structures, we performed additional DALI searches submitting individual domains.

We found that both the N- and C-terminal domains of pORF23 showed structural homology to regions of the CVSC/ PVAT protein pUL17 of herpes simplex virus type1 (PDB 6ODM chain C – Movie S3) (*10*). The N-terminal domain includes a C2 domain (aa 80-168) that most closely aligned to that of Phosphatidylinositol 4-Phosphate 3-Kinase C2α (PDB 7BI2 chain A aa 303-458 Z=5.7 RMSD=2.7) (*37*). This motif is however also present in pUL17 (PDB 6ODM chain C aa 22-150,383-403 Z=2.3, RMSD=4.2). The C-terminal domain of pORF23 (aa 263-349) also aligned to a separate region of pUL17 (PDB 6ODM chain C aa 585-671 Z=2.9, RMSD=3.6). Although the individual alignments have low Z-scores, together they point to a conservation of structure. The presence of two topologically similar domains in pORF23 and pUL17 is less parsimoniously explained by convergent evolution than similarity confined to a single domain. Convergence typically yields localized resemblance driven by functional constraints, whereas the replication of complex β-sheet topology across multiple domains is statistically less probable. This pattern therefore more strongly supports inheritance from a common ancestral protein followed by divergence, rather than independent structural convergence. The functional roles of pORF23 and pUL17 as the foundation of the PVAT in their respective viruses further supports the suggestion that pORF23 and pUL17 are orthologous proteins.

### pORF67 comprises multiple putative macrodomains

As well as the homology between the structures of pORF23 and the orthoherpesvirus CVSC/PVAT protein pUL17, compelling hits were found for pORF67 (Figure 6, Movie S3). We found four distinct domains that were readily aligned to known structures of macrodomains of multiple species. The most confident alignments for a pORF67 domain with known macrodomains were at aa 946-1146. This region aligns well to PARP9 macrodomain 2 (PDB 9QYH aa 310-490, Z=8.8, RMSD=3.0 Å), PARP14 macrodomain 3 (PDB 5QI8 aa 1207-1388, Z=8.4 RMSD=2.8 Å), and to SARS-CoV2 Mac1 domain of nsp3 (PDB 9GUB Z=5.3, RMSD=3.1 Å) (*38*). Two domains were identified that match macrodomains 1 and 2 of PARP14 at pORF67 aa 62-248 (PDB 3VFQ aa 791-978, Z=3.9, RMSD=3.4 Å) and 330-458 (PDB 3VFQ aa 1005-1191, Z=6.5, RMSD=3.6 Å) respectively (*39*). Finally a fourth domain at pORF67 aa 1285-1446 was found to align to the macrodomain of the protozoal parasite *Trypanosoma cruzi* the causative agent of Chagas disease (PDB 5FSZ aa 163-257, Z=4.3, RMSD=3.0 Å) (*40*).

**Figure 6:**
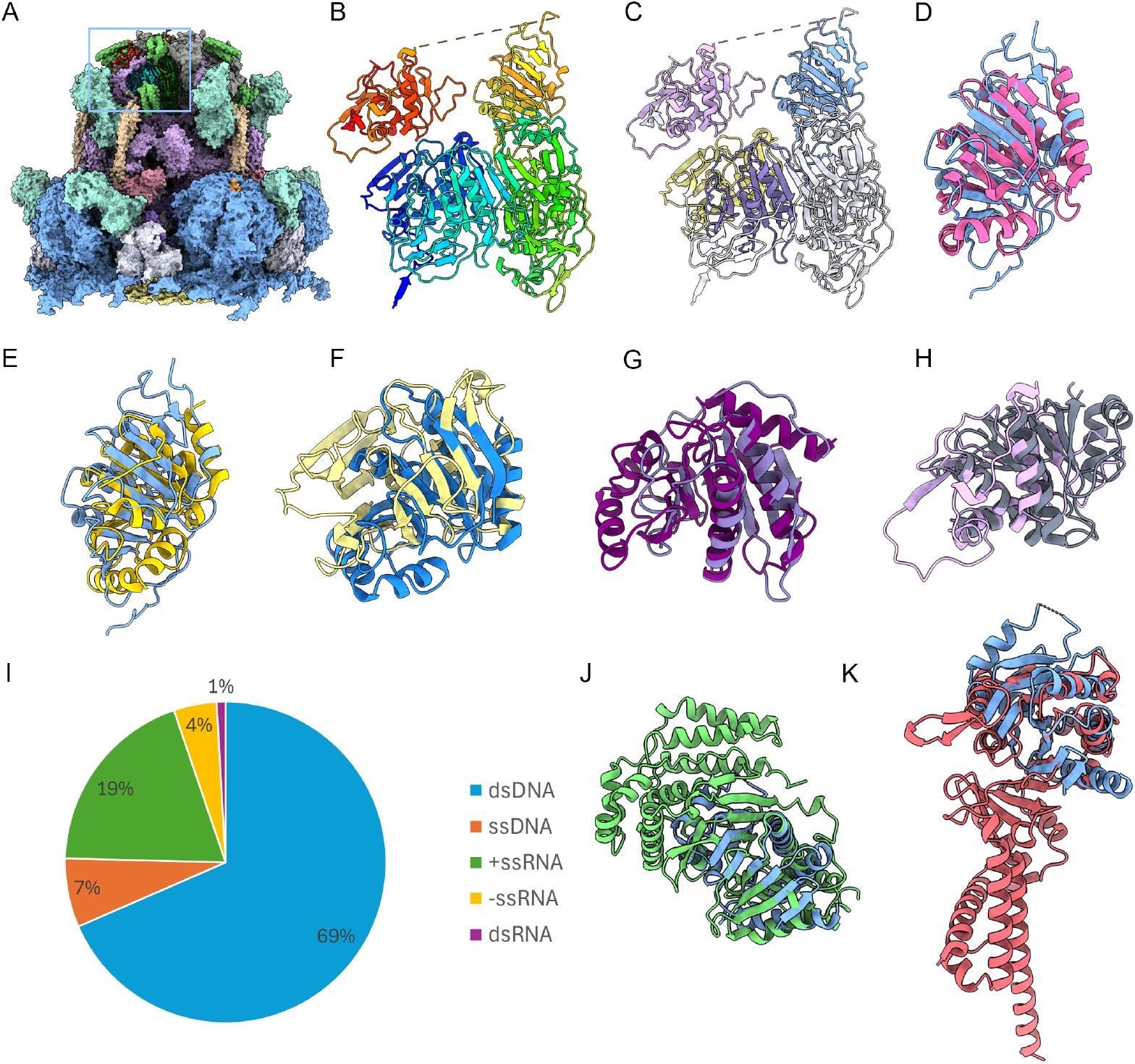
pORF67 comprises multiple macrodomain-like folds. (A) Side view of the PVAT complex coloured according to the scheme presented in figure 5. A pORF67 monomer is shown as a rainbow-coloured ribbon diagram, highlighted by a blue box. (B-C) The structure of pORF67 is shown in the same orientation as (A) in rainbow colour (B) and coloured to highlight individual domains (C). (D) pORF67 aa 946-1146 (blue) aligned to PARP9 macrodomain 2 (PDB 9QYH – pink). (E) pORF67 aa 946-1146 (blue) aligned to the SARS-CoV2 Mac1 domain (PDB 9GUB – yellow (*38*)). (F) pORF67 aa 62-258 (yellow) aligned to PARP14 macrodomain 1 (PDB 3VFQ – blue (*39*)). (G) pORF67 aa 330-458 (light mauve) aligned to PARP14 Macrodomain 2 (PDB 3VFQ – dark purple (*39*)). (H) pORF67 aa 1285-1446 (pink) aligned to the Trypanosoma cruzi macrodomain (PDB 5FSZ – grey (*40*)). (I) To explore the extent of macrodomain-like folds in the virosphere we submitted the PARP9 and PARP14 macrodomain structures to a DALI search of the Viro3D database. The hits were categorised according to the genome content of each virus identified. (J) The predicted structure of pORF105, the Ac114 protein of Choristoneura murinana alphabaculovirus (green) and (K) adenovirus D32 pIVa2 (red) aligns well to PARP9 macrodomain 2 (blue).

Originally identified as ‘X-domains’ in coronaviruses, alphaviruses and rubiviruses (*41*), macrodomains are named after the Macro-H2A histone protein C-terminal domain (*42*) and play a critical role in regulating gene expression by binding ADP-ribose moieties (*43*). Mono- and poly-ADP-ribosylation (MARylation and PARylation) are critical modifications of proteins and nucleic acids, that are added by several classes of cellular proteins including poly-ADP-ribose polymerases (PARPs). PARPs regulate a wide range of biological functions, including DNA repair, chromatin remodelling, transcriptional regulation and immune responses. Humans express 17 PARPs, three of which incorporate macrodomains. Importantly some macrodomains including several encoded by positive-sense RNA viruses catalyse the hydrolysis of ADP-ribose moieties and are thought thereby to down-regulate host innate immunity (*44-46*). Since macrodomains often occur as repeating tandem units within a single protein (such as coronavirus nsp3). The presence of four such domains in pORF67 strongly suggests a functional arrangement, however, upon inspection of the four pORF67 macrodomains we did not find identifiable MAR binding clefts at the expected locations when aligned with their respective PARP homologues. Thus, experimental confirmation of ligand-binding or enzymatic properties is required before we ascribe a function to these domains.

For the remaining IcHV-1 proteins identified, pORF24, 65 and 66 no convincing homology was found to known protein structures.

### Macrodomains may be more widespread among viruses than previously suspected

Macrodomains have been identified and widely studied in several RNA viruses. In contrast, their presence in DNA viruses is less well characterised, although they have been reported in members of the *Iridoviridae (47*) and *Poxviridae (48*). Our findings prompted us to explore whether macrodomains might be found in other virus families. Since experimentally determined structure data are limited, we integrated the recently published Viro3D database of predicted protein structures into the DALI search (*30*). In our analysis of the IcHV-1 pORF67 structure, the macrodomains that gave the strongest hits were PDB 9QYH, PARP9 and PDB 3VFQ, PARP14. When we used these data to search Viro3D we found predicted macrodomain-like folds in both DNA and RNA viruses (Figure 6I). The analysis confirmed their widespread presence in members of the *Coronaviridae, Hepeviridae, Iridoviridae, Matonaviridae* (rubiviruses) *Poxviridae*, and *Togaviridae*, but also identified putative macrodomains in members of the *Adenoviridae, Baculoviridae, Herpesviridae, Nimaviridae, Peribunyaviridae*, and *Reoviridae*. These hits all had acceptable Z-scores (cutoff 5) and RMSD measurements. For example, the predicted structure of pORF105, the Ac114 protein of Choristoneura murinana alphabaculovirus gave a Z-score of 10.2 and RMSD of 3.1 Å (Figure 6J), and the adenovirus D32 pIVa2 protein gave a Z-score of 5.5 and RMSD of 3.3 Å (Figure 6K). Definitive identification of functional macrodomains would require experimental testing of MAR binding and ADP-ribosyl hydrolase activity, but these data suggest a broader distribution of this fold across the virosphere.

## Discussion

The alloherpesvirus IcHV-1 is highly diverged from the well-characterised orthoherpesviruses that infect humans. The only hallmark gene that can be used for sequence-based assembly of a phylogeny of the *Herpesvirales* is the terminase gene. IcHV-1 ORFs were not able to be annotated confidently without the application of proteomic analysis of purified capsids and virions, and even then, this proved challenging (*22, 23*). We used cryo-EM combined with the ground-breaking machine-learning tool for atomic model construction ModelAngelo, to determine the high-resolution structures for both the immature capsid and mature virion of IcHV-1. The application of symmetry-expansion and focussed classification allowed us to relax the full icosahedral symmetry that was initially employed, to reconstruct the 5-fold symmetric capsid and virion to reveal their portal vertices. This led to our surprising discovery that, unlike orthoherpesvirus structures, the IcHV-1 virion has a substantial ‘tail’ or portal-vertex associated tegument. To achieve high-resolution cryo-EM maps required careful particle subtraction and reconstruction of individual capsomeres and portal-vertex components as well as masked local-refinement. This led to calculation of maps that ranged in resolution from 3 to 4.5 Å. Despite the modest resolution of some maps, we were able to unambiguously assign identities and build models for the majority of viral capsid and portal-vertex associated proteins captured by the reconstruction process. This was entirely owing to the surprising capacity of ModelAngelo to build high-quality models into cryo-EM maps at poorer than 4 Å resolution, even in the absence of supporting sequence data. By submitting built sequences to BLAST, or using the integrated HMMER search tool, we identified ten structural proteins: the major capsid protein pORF39, the triplex proteins pORF53 and 27, the inner tegument protein pORF65, the hexon channel protein pORF42, the portal protein pORF37, and four further PVAT proteins, pORF23, 24, 66 and 67.

Whereas the immature capsid portal-vertex presented only as an opening in the capsid shell with the poorly constrained portal at the interior, the virion portal-vertex was extensive and elaborate – comprising a total of 67 polypeptide chains, excluding the surrounding hexons and triplexes, for which we have modelled a total of 27,282 amino acid residues (including the 5- and 12-fold symmetry). Combining these data with the icosahedral components of the virion allowed us to assemble a complete model comprising some two million amino-acid residues and more than 20 million atoms (Figure 7).

**Figure 7:**
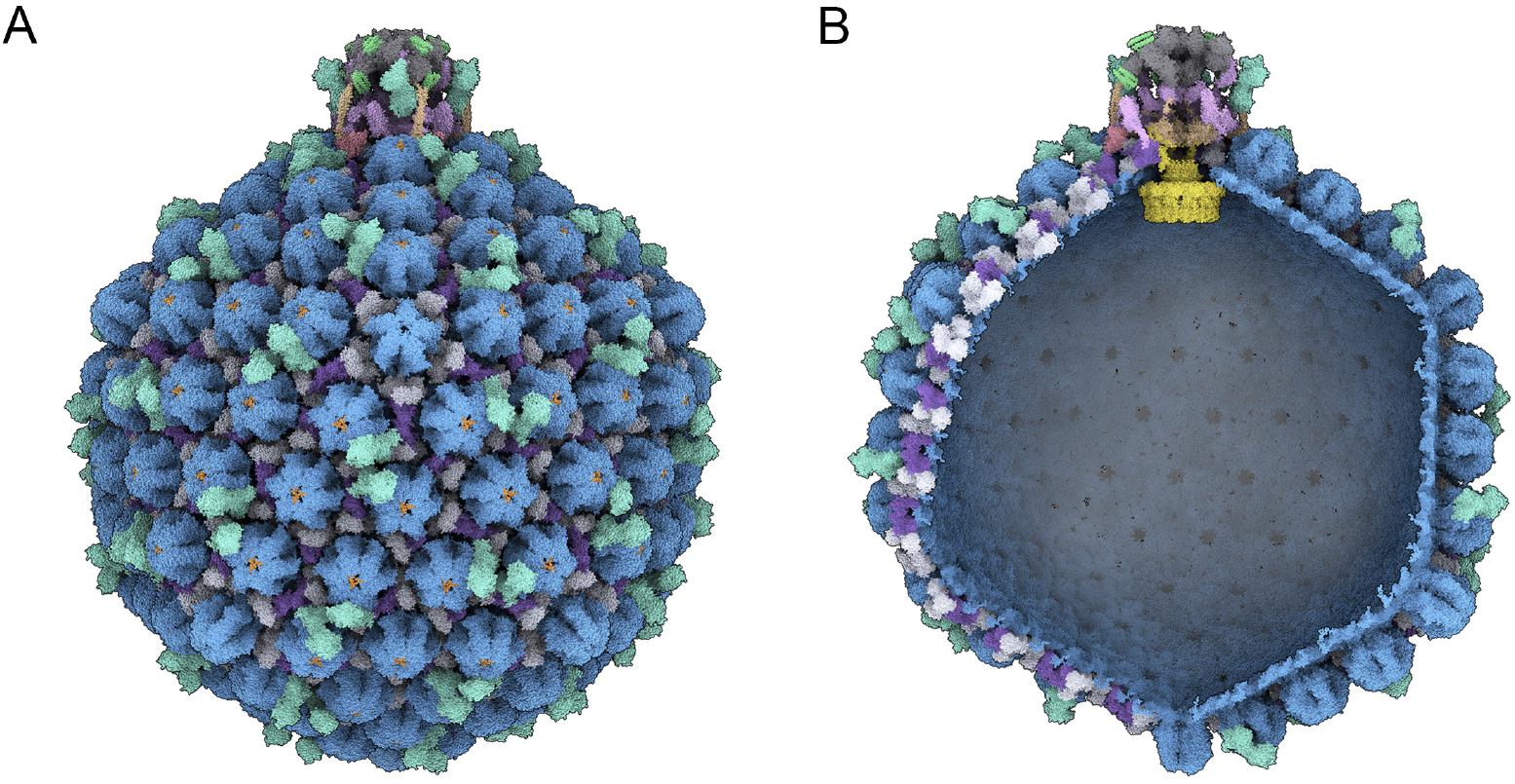
A composite model of the IcHV-1 virion. (A) All atom representation (spheres) of the virion structure including the complete capsid shell and PVAT, coloured according to the scheme used in figures 2 and 5. (B) Cutaway view to reveal the atomic model of the symmetry mismatched portal and interior of the capsid and PVAT.

Our data revealed a high degree of structural conservation in the core capsid proteins pORF39, 53, 27 and 37. Structures of their orthologues are published for several human-infecting orthoherpesviruses. The MCP presented a similar topology to those described previously, with the one notable feature of having a C-terminal arm that inserts into the middle section of a neighbouring protomer to stabilise hexon assemblies; a domain insertion that is not present in the penton. The triplex proteins pORF53 and pORF27 also show an overall conserved structure, with some interesting observations at the virion portal-vertex, where the Ta triplex is rotated compared to that at penton vertices, to engage the turret/tentacle-helices of the portal pORF37. A bridge formed of the C-terminal portion of pORF53 extends towards the upper domain of the MCP at position P3 of the peri-portal P-hexon to support the assembly of the large PVAT. Finally, the portal protein pORF37 shows considerable similarity to published herpesvirus and phage portal proteins, forming a C12 symmetric wheel shaped assembly having the same turret, clip, stem, wing, β-hairpin/tunnel-loop, and wall domains. The configuration of the wall and wing domains give rise to a different overall morphology when compared to known orthoherpesvirus portal complexes. Interestingly in our structure the tunnel-loop is not stabilised by an interaction with the terminal DNA that extends through the portal. Rather, constrictions in the distal PVAT seem to constrain the packaged genome from premature release. We also noted an interesting rearrangement of the portal wing domain comparing the capsid and virion structures, whereby in the virion portal, the N-terminal sequence engages neighbouring protomers via β-augmentation possibly to stabilise the assembly.

The extensive PVAT we have noted for IcHV-1 came as a surprise and led to the identification and modelling of several novel proteins. However, given the considerable internal pressure exerted by the packaged genome within orthoherpesviruses (*49*), we should perhaps be more surprised by the apparent lack of an extensive tail in e.g. herpes simplex virus (*10, 11*). In that virus, interactions between the terminal packaged DNA and the portal tunnel-loop, and the poorly resolved double-tiered PVAT in which ten copies of pUL25 cap the decameric tentacle helices, are the only apparent means of genome retention. In our cryo-EM map of the IcHV-1 portal vertex, several constrictions of the DNA translocation tunnel involving pORF24, 66 and 67 along with the capping of the PVAT by pORF67 all appear to act to hold the genome in place.

While many of the PVAT proteins appeared to present novel folds, we observed that pORF67 includes four domains having macrodomain-like topology. Herpesvirus tegument proteins sculpt the cellular environment to favour viral replication. In several RNA viruses macrodomains have been shown to modulate host immune responses by antagonising PARP activity. PARP proteins regulate transcription by polymerisation and in some cases removal of poly-ADP ribose, PARP 9 in particular plays a key role in host immunity, triggering type I interferon production in response to viral infection (*50*). Viral antagonism of innate immunity is critical to establish a productive infection. Members of the *Coronaviridae* including severe acute respiratory syndrome (SARS) viruses 1 and 2, achieve this by encoding macrodomains in nsp3. One of these – the Mac1 domain – has been shown to have ADP-ribosyl hydrolase activity *in vitro*, while mutation leads to an attenuation phenotype that rapidly triggers innate immunity upon infection (*51, 52*). It is tempting to surmise then that one possible role for pORF67 is in suppressing the host immune response to infection. The apparent lack of an ADP-ribose binding cleft in the observed macrodomains might suggest otherwise, however. Two further nsp3 macrodomains that make up the SARS Unique Domain (SUD), SUD-N and SUD-M, do not bind ADP-ribose and instead bind to DNA and RNA oligonucleotides that form guanine-quadruplexes (G4) (*53*). G4s are enriched in regulatory regions of eukaryotic genomes as well as in RNA and DNA viruses. Mutation of the SUD macrodomains impacts on viral replication and transcription. Thus, direct interactions with regulatory elements in either the host or viral genome may be a plausible function for pORF67 macrodomains. Finally, coronavirus nsp3 has also been shown to be a critical component of the crown complex, a pore-like assembly that is formed in double-membrane vesicles (DMVs) (*54*). DMVs are virus induced cellular compartments that sequester the viral replicative machinery (*55*). Spanning the double-membrane of these replication organelles, crown complexes are RNA translocation pores that extrude newly synthesised viral genomes to the cytosol for encapsidation. Further parallel’s might be drawn between nsp3 and pORF67 indicating a potential functional role in genome translocation. Further work is warranted to explore the role of pORF67 in IcHV-1 pathogenesis and may lead to the identification of new targets for therapeutic intervention or attenuation strategies for vaccine production.

Our unexpected discovery of macrodomain-like folds in IcHV-1 pORF67 led us to explore the extent to which these features might be found in other viruses. While the presence of macrodomains in positive-sense RNA containing viruses such as coronaviruses and alphaviruses is quite well documented, there is a paucity of data pertaining to their presence and function in DNA viruses. Consultation of the literature uncovered limited studies, identifying a putative macrodomain in salmon gill pox virus (*48*) and a functional macrodomain in infectious spleen and kidney necrosis virus (*47*). Interestingly, both viruses are also notable aquaculture pathogens. To systematically explore the prevalence of this protein domain across the virosphere, we added the viro3D database to the search capabilities of DALI, a widely used tool for comparing the 3D structures of proteins (*30, 36*). Searching this large database of predicted protein structures against the known structures of PARP9 and PARP14 macrodomains returned an abundance of previously unannotated putative viral macrodomains. Confirmation of these proteins as *bona-fide* macrodomains will require individual biochemical and structural characterisation. The data do however suggest that this fold may be found in a broader spectrum of viral families than previously thought.

The potential to search large databases of predicted protein structures represents an exciting new research tool – AI based function discovery. Conservation of protein folds persists over far longer evolutionary periods than sequence, as evidenced by the conservation of core capsid proteins in this study. Thus, this new capability may enable researchers to ascribe functions to viral proteins and perhaps develop new approaches to intervention. For example, highly druggable known protein structures could be used to search the databases of viral protein predicted structures to identify viral proteins that may be susceptible to known pharmacophores, providing new testable hypotheses for antiviral development.

Overall, our study has provided a detailed structural characterisation of IcHV-1, a notable aquaculture pathogen and evolutionarily distant herpesvirus. Our analysis yielded several surprising insights, including the presence of a substantial portal-vertex associated tail and a large tegument protein that comprises several macrodomain-like folds. Our findings led us to establish a new capability within the DALI search tool that enables searching of several databases of predicted viral protein structures (Viro3D, VAD, BFVD - (*30, 56, 57*)). This allowed us to identify putative macrodomains in the proteomes of many virus families previously not described as having them. Taken together, our analysis has revealed both structural conservation and novelty in the *alloherpesviridae*, uncovered a potential target for the development of antivirals or vaccines against this family, and established a new tool to discover functional protein domains by searching a large database of predicted viral protein structures.

## Methods

### Phylogenetic analysis

The maximum likelihood phylogeny was inferred using MEGA X (*58, 59*) with a published amino acid replacement model (*60*). The tree with the highest log likelihood (−19387.21) under the LG+G+I model is shown, with bootstrap values (from 100 runs) at the nodes. All positions with less than 90% site coverage were eliminated, leaving 575 positions in the final dataset.

### Virus culture

IcHV-1 strain Auburn 1 (ATCC VR-665) was propagated in brown bullhead (BB) cells (ATCC CCL-59) in 30 T-175 flasks. Cells were maintained in MEM Eagles medium containing Earle’s salts, L-glutamine and sodium bicarbonate, supplemented with 10% foetal calf serum and 1% non-essential amino acids, and incubated at 28°C. After 3-days the cell-culture medium was decanted and virions were purified. Infected cells were retained separately, and capsids were purified from the fractionated nuclei.

### Virion purification

The culture medium was clarified by centrifugation at 1500*g* for 30 minutes at 4°C. Virus particles in the clarified medium were pelleted by centrifugation (15000*g* for 2 hr at 4 °C). The pellet was gently resuspended in 2 ml culture medium, layered on a 35 ml pre-formed gradient of 5-15% Ficoll 400 and centrifuged using a swing-out rotor (26000*g* for 2 hr at 4 °C). The virion band was withdrawn by side puncture. Finally, the virions were pelleted by centrifugation (72000*g* for 2 hr at 4 °C), and gently resuspended in 100ml phosphate-buffered saline (PBS)

### Capsid purification

Infected cells were pelleted at 1500*g* for 10 minutes, resuspended in Tris-buffered saline (NTE) containing 10% IGEPAL detergent and incubated on ice for 10 minutes. Cell nuclei were collected by centrifugation at 1500*g* for 10 minutes, resuspended in PBS and lysed by sonication. Debris were removed by centrifugation (1500*g* for 10 minutes) and the supernatant was then layered onto a cushion of 40% sucrose in NTE and centrifuged at 72000*g* for 1hr. The pellet was resuspended in 1ml PBS/IGEPAL, DNAse-treated, sonicated and clarified by centrifugation at 1500*g* for 10 minutes. The supernatant was loaded onto a 5-40% gradient of sucrose in NTE and centrifuged at 72000*g* for 1hr. The upper band corresponding to empty capsids was removed by side-puncture, and capsids were pelleted by centrifugation at 72000*g* for 1 hr at 4 °C and then gently resuspended in 100ml PBS.

### Cryo-electron microscopy

IcHV-1 capsids and virions were prepared for cryo-EM by plunge freezing into liquid-nitrogen cooled liquid ethane using a Thermo Fisher Vitrobot Mark IV. Briefly, 3 ml of virion or capsid suspension was pipetted onto a freshly glow-discharged Quantifoil (R2/2) holey carbon coated TEM grid, at 100% humidity and at 4°C. The excess liquid was blotted away with a blot time of 3 seconds. The grid was then plunged rapidly into liquid ethane and stored under liquid nitrogen.

Grids were imaged at the Scottish Centre for Macromolecular Imaging in a JEOL CryoARM 300 automated cryomicroscope equipped with a Direct Electron Apollo detector. All automated data collection was performed using the SerialEM software package (*61*). Micrograph movies were collected at 60,000’ magnification (corresponding to 1.05Å per physical pixel) with the energy filter slit width set to 20 eV. The Apollo camera integrates counted electron events at 60 frames/s and applies event centroiding by default to capture 2’ super-resolution images. Exposures of 2s were captured, integrating every three counted frames such that 40 super-resolution frames were saved per micrograph movie. The total electron dose per micrograph movie was ∼60e/Å2. Micrograph movies were saved as un-corrected frames in MRC format and compressed to LZW-TIFF format before processing.

### Image processing and three-dimensional image reconstruction

Capsid and virion data were processed using a common strategy to resolve symmetrically ordered and asymmetric features using the RELION-5 software package (*62, 63*) running on a high-performance GPU computing cluster. Micrograph movies were motion-corrected using the RELION implementation of the MotionCor2 algorithm (*64*). The defocus and objective lens astigmatism was then estimated using CTFfind4 (*65*). A subset of particles was selected manually and used to train a TOPAZ automatic picking model (*66*) that was then used to pick the full dataset of particles. Two-dimensional classification was used to curate the particle dataset and remove incorrect picks. *Ab-initio* starting models were generated using the stochastic gradient descent algorithm and imposing I2 symmetry (*67*). Particle images were extracted at a pixel size of 1.05Å in boxes of 1600’1600 pixels and three-dimensional refinement was then used to determine accurate orientations and origins for each particle image using I2 and subsequently I4 symmetry which better enabled resolution of symmetry-mismatched features at the five-fold portal vertex.

To achieve high-resolution maps suited to construction of atomic models, a process of symmetry expansion and particle subtraction was used to reconstruct the four unique capsomeres: penton, P-hexon, E-hexon and C-hexon and their surrounding triplexes. A cylindrical mask was created in ChimeraX (*68*) with a diameter and length of 340Å. The mask was then positioned to cover each capsomere within the asymmetric unit and resampled into a 1600’1600’1600 voxel box to produce four masks. Particle orientations were symmetry-expanded to derive 60 views for each particle reflecting the 60-fold redundancy of the icosahedral capsid. Particle subtraction was then used to extract images of each capsomere in every capsid or virion in a 340’340 pixel box. These images were used to calculate a starting map and the orientations and origins of each capsomere image were then subjected to local C1 refinement until they converged.

To calculate 3D reconstructions of the capsid and virion portal vertices and symmetry-mismatched portal complexes, the orientations of each particle image were refined with I4 symmetry imposed. This places the icosahedral five-fold axis on the z-axis of the reconstruction, thus simplifying the process of resolving the symmetry mismatch. Particles were extracted with 5’ binning, and their orientations were symmetry-expanded. A cylindrical mask was applied to the five-fold symmetry axis congruent with the z-axis of the 3D reconstruction. Focussed classification was performed without orientation refinement and with a tau value of 20. Portal-vertex classes were selected, and particle images were then extracted from the unbinned particle images by particle-subtraction in 600’600 pixel boxes. Portal vertex images were then reconstructed. To calculate a high-resolution map of the five-fold symmetric features, duplicate particle images were removed, and the orientations and origins were subjected to local-refinement with C5 symmetry imposed. To improve the resolution of the distal features of the virion PVAT a mask was created by erasing the capsid shell density leaving only exterior PVAT density. This was low-pass filtered and dilated, and a soft edge was applied. Masked local refinement was then performed with C5 symmetry. To resolve the structure of the symmetry mismatched C12 portal, a mask was created by deleting the capsid shell density in the capsid portal-vertex reconstruction, leaving only the incoherently averaged portal density. This was low-pass filtered and dilated, and a soft edge applied. Masked focussed classification of the portal-vertex images was then performed without refinement, with a tau value of 20 and with C12 symmetry applied. Classes that showed clear portal density were selected, duplicate images were removed and the remaining images were subjected to local refinement. In each case the map that achieved the highest resolution was used for model building.

### Model building

To confirm the identities of each protein and construct atomic models we used the ModelAngelo programme within Relion (*25*). Each map was submitted to the programme without sequence data. The resulting model was then inspected in ChimeraX, larger model fragments were selected and their sequences were extracted and submitted to the NCBI BLASTP tool (*26*). When a sequence was matched to an IcHV-1 ORF, the sequence of that protein was used subsequently for a second round of ModelAngelo. In this round of model building a single FASTA file containing the combined protein sequences for all identified IcHV-1 structural proteins was supplied. Models were then either revised in Coot (*69*) and ChimeraX, to combine discontinuous protein chains, or predicted models from the viro3D database (*70*) were aligned to the ModelAngelo output using Matchmaker in ChimeraX and then manually remodelled to fit the map density using the ISOLDE plugin (*71*). All models were refined in ISOLDE followed by automated refinement in Phenix (*72*) using refinement parameters generated by ISOLDE. Maps and models were visualised in ChimeraX (*68*).

### Structure-based searches for homologous and orthologous proteins

To identify conserved folds in newly solved protein structures we used the DALI search tool to search against the protein data bank, using whole proteins and individual protein domains (http://ekhidna2.biocenter.helsinki.fi/dali) (*73, 74*). To enable searching against databases of predicted viral proteins, a new functionality was added to the DALI search tool incorporating systematic searching of Viro3D (*30*) (https://viro3d.cvr.gla.ac.uk) and the Viral Alphafold Database (*56*) (VAD - https://vad.atkinson-lab.com) (*75*), and iterative searching of the Big Fantastic Virus Database (BFVD) (*57, 76*).

## Acknowledgements

This work was supported by the United Kingdom Medical Research Council (MC_UU_12014/6, MC_UU_12014/7, MC_ UU_00034/1) and the Research Council of Finland (357350). We acknowledge the Scottish Centre for Macromolecular Imaging (SCMI) for access to cryo-EM instrumentation, funded by the United Kingdom Medical Research Council (MC_PC_17135, MC_UU_00034/7, MR/X011879/1) and SFC (H17007).

The MRC – University of Glasgow Centre for Virus Research uses the CRediT contributor role taxonomy. AM – Formal analysis, Investigation, Methodology, Validation, Visualisation, Writing – Original Draft. FP – Data Curation, Formal analysis, Investigation, Methodology, Validation. MM – Investigation. JS – Investigation, Resources. LH – Formal analysis, Methodology, Software, Validation, Writing – review and editing. JG – Conceptualisation, Methodology, Resources, Writing – review and editing. AJD – Data curation, Formal analysis, Methodology, Writing – review and editing. FJR Conceptualisation, Funding acquisition, Investigation, Writing – review and editing. DB – Conceptualisation, Data curation, Formal analysis, Funding acquisition, Investigation, Methodology, Project administration, Resources, Supervision, Validation, Visualisation, Writing – original draft.

## References

1. D. Gatherer et al., ICTV Virus Taxonomy Profile: Herpesviridae 2021. J Gen Virol 102, (2021).

2. D. J. McGeoch, S. Cook, A. Dolan, F. E. Jamieson, E. A. Telford, Molecular phylogeny and evolutionary timescale for the family of mammalian herpesviruses. J Mol Biol 247, 443–458 (1995).

3. D. J. McGeoch, F. J. Rixon, A. J. Davison, Topics in herpesvirus genomics and evolution. Virus Res 117, 90–104 (2006).

4. Z. H. Zhou et al., Seeing the herpesvirus capsid at 8.5 A. Science 288, 877–880 (2000).

5. D. H. Bamford, J. M. Grimes, D. I. Stuart, What does structure tell us about virus evolution? Curr Opin Struct Biol 15, 655–663 (2005).

6. A. J. Davison, D. Bhella, in Human Herpesviruses: Biology, Therapy, and Immunoprophylaxis, A. Arvin et al., Eds. (Cambridge, 2007).

7. T. Aoki et al., Genome sequences of three koi herpesvirus isolates representing the expanding distribution of an emerging disease threatening koi and common carp worldwide. J Virol 81, 5058–5065 (2007).

8. X. Dai, Z. H. Zhou, Structure of the herpes simplex virus 1 capsid with associated tegument protein complexes. Science 360, (2018).

9. X. Yu, J. Jih, J. Jiang, Z. H. Zhou, Atomic structure of the human cytomegalovirus capsid with its securing tegument layer of pp150. Science 356, (2017).

10. Y. T. Liu, J. Jih, X. Dai, G. Q. Bi, Z. H. Zhou, Cryo-EM structures of herpes simplex virus type 1 portal vertex and packaged genome. Nature 570, 257–261 (2019).

11. M. McElwee, S. Vijayakrishnan, F. Rixon, D. Bhella, Structure of the herpes simplex virus portal-vertex. PLoS Biol 16, e2006191 (2018).

12. Y. Yang et al., Architecture of the herpesvirus genome-packaging complex and implications for DNA translocation. Protein Cell 11, 339–351 (2020).

13. D. Bhella, F. J. Rixon, D. J. Dargan, Cryomicroscopy of human cytomegalovirus virions reveals more densely packed genomic DNA than in herpes simplex virus type 1. J Mol Biol 295, 155–161 (2000).

14. J. Jih, Y. T. Liu, W. Liu, Z. H. Zhou, The incredible bulk: Human cytomegalovirus tegument architectures uncovered by AI-empowered cryo-EM. Sci Adv 10, eadj1640 (2024).

15. W. H. Fan et al., The large tegument protein pUL36 is essential for formation of the capsid vertex-specific component at the capsid-tegument interface of herpes simplex virus 1. J Virol 89, 1502–1511 (2015).

16. S. L. Oliver et al., Cryogenic Electron Tomography Redefines Herpesvirus Capsid Assembly Intermediates Inside the Cell Nucleus. bioRxiv, 2025.2006.2027.661840 (2025).

17. S. Vijayakrishnan, M. McElwee, C. Loney, F. Rixon, D. Bhella, In situ structure of virus capsids within cell nuclei by correlative light and cryo-electron tomography. Sci Rep 10, 17596 (2020).

18. W. W. Newcomb et al., The Primary Enveloped Virion of Herpes Simplex Virus 1: Its Role in Nuclear Egress. mBio 8, (2017).

19. M. H. Bergeman et al., Individual herpes simplex virus 1 (HSV-1) particles exit by exocytosis and accumulate at preferential egress sites. J Virol 98, e0178523 (2024).

20. N. N. Fijan, AN ACUTE VIRAL DISEASE OF CHANNEL CATFISH. Fish and Wildlife Service Technical Paper 43, (1970).

21. K. Wolf, R. W. Darlington, Channel catfish virus: a new herpesvirus of ictalurid fish. J Virol 8, 525–533 (1971).

22. A. J. Davison, Channel catfish virus: a new type of herpesvirus. Virology 186, 9–14 (1992).

23. A. J. Davison, M. D. Davison, Identification of structural proteins of channel catfish virus by mass spectrometry. Virology 206, 1035–1043 (1995).

24. F. P. Booy, B. L. Trus, A. J. Davison, A. C. Steven, The capsid architecture of channel catfish virus, an evolutionarily distant herpesvirus, is largely conserved in the absence of discernible sequence homology with herpes simplex virus. Virology 215, 134–141 (1996).

25. K. Jamali et al., Automated model building and protein identification in cryo-EM maps. Nature 628, 450–457 (2024).

26. S. F. Altschul, W. Gish, W. Miller, E. W. Myers, D. J. Lipman, Basic local alignment search tool. J Mol Biol 215, 403–410 (1990).

27. J. Mistry, R. D. Finn, S. R. Eddy, A. Bateman, M. Punta, Challenges in homology search: HMMER3 and convergent evolution of coiled-coil regions. Nucleic Acids Res 41, e121 (2013).

28. S. R. Eddy, Accelerated Profile HMM Searches. PLoS Comput Biol 7, e1002195 (2011).

29. M. Mirdita et al., ColabFold: making protein folding accessible to all. Nature Methods 19, 679–682 (2022).

30. U. Litvin et al., Viro3D: a comprehensive database of virus protein structure predictions. Mol Syst Biol 21, 1599–1617 (2025).

31. J. Sun et al., Cryo-EM structure of the varicella-zoster virus A-capsid. Nat Commun 11, 4795 (2020).

32. Z. Li et al., CryoEM structure of the tegumented capsid of Epstein-Barr virus. Cell Res 30, 873–884 (2020).

33. A. Cuervo et al., Structures of T7 bacteriophage portal and tail suggest a viral DNA retention and ejection mechanism. Nat Commun 10, 3746 (2019).

34. M. van Kempen et al., Fast and accurate protein structure search with Foldseek. Nat Biotechnol 42, 243–246 (2024).

35. Z. Li, J. Pang, L. Dong, X. Yu, Structural basis for genome packaging, retention, and ejection in human cytomegalovirus. Nat Commun 12, 4538 (2021).

36. L. Holm, A. Laiho, P. Toronen, M. Salgado, DALI shines a light on remote homologs: One hundred discoveries. Protein Sci 32, e4519 (2023).

37. W. T. Lo et al., Structural basis of phosphatidylinositol 3-kinase C2alpha function. Nat Struct Mol Biol 29, 218-228 (2022).

38. J. J. Pfannenstiel et al., Identification of a series of pyrrolo-pyrimidine-based SARS-CoV-2 Mac1 inhibitors that repress coronavirus replication. mBio 16, e0386524 (2025).

39. A. H. Forst et al., Recognition of mono-ADP-ribosylated ARTD10 substrates by ARTD8 macrodomains. Structure 21, 462–475 (2013).

40. T. Haikarainen, L. Lehtio, Proximal ADP-ribose Hydrolysis in Trypanosomatids is Catalyzed by a Macrodomain. Sci Rep 6, 24213 (2016).

41. H. J. Lee et al., The complete sequence (22 kilobases) of murine coronavirus gene 1 encoding the putative proteases and RNA polymerase. Virology 180, 567–582 (1991).

42. L. Aravind, The WWE domain: a common interaction module in protein ubiquitination and ADP ribosylation. Trends Biochem Sci 26, 273–275 (2001).

43. G. I. Karras et al., The macro domain is an ADP-ribose binding module. EMBO J 24, 1911–1920 (2005).

44. B. Luscher et al., ADP-ribosyltransferases, an update on function and nomenclature. FEBS J 289, 7399–7410 (2022).

45. C. Li et al., Viral Macro Domains Reverse Protein ADP-Ribosylation. J Virol 90, 8478–8486 (2016).

46. L. Eckei et al., The conserved macrodomains of the non-structural proteins of Chikungunya virus and other pathogenic positive strand RNA viruses function as mono-ADP-ribosylhydrolases. Sci Rep 7, 41746 (2017).

47. J. G. M. Rack et al., Viral macrodomains: a structural and evolutionary assessment of the pharmacological potential. Open Biol 10, 200237 (2020).

48. M. C. Gjessing et al., Salmon Gill Poxvirus, the Deepest Representative of the Chordopoxvirinae. J Virol 89, 9348–9367 (2015).

49. D. W. Bauer, J. B. Huffman, F. L. Homa, A. Evilevitch, Herpes virus genome, the pressure is on. J Am Chem Soc 135, 11216–11221 (2013).

50. J. Xing et al., Identification of poly(ADP-ribose) polymerase 9 (PARP9) as a noncanonical sensor for RNA virus in dendritic cells. Nat Commun 12, 2681 (2021).

51. A. R. Fehr et al., The Conserved Coronavirus Macrodomain Promotes Virulence and Suppresses the Innate Immune Response during Severe Acute Respiratory Syndrome Coronavirus Infection. mBio 7, (2016).

52. K. K. Eriksson, L. Cervantes-Barragan, B. Ludewig, V. Thiel, Mouse hepatitis virus liver pathology is dependent on ADP-ribose-1’’-phosphatase, a viral function conserved in the alpha-like supergroup. J Virol 82, 12325–12334 (2008).

53. J. Tan et al., The SARS-unique domain (SUD) of SARS coronavirus contains two macrodomains that bind G-quadruplexes. PLoS Pathog 5, e1000428 (2009).

54. Y. Huang et al., Molecular architecture of coronavirus double-membrane vesicle pore complex. Nature 633, 224–231 (2024).

55. G. Wolff et al., A molecular pore spans the double membrane of the coronavirus replication organelle. Science 369, 1395–1398 (2020).

56. R. Odai et al., The Viral AlphaFold Database of monomers and homodimers reveals conserved protein folds in viruses of bacteria, archaea, and eukaryotes. Sci Adv 11, eadz8560 (2025).

57. R. S. Kim, E. Levy Karin, M. Mirdita, R. Chikhi, M. Steinegger, BFVD-a large repository of predicted viral protein structures. Nucleic Acids Res 53, D340–D347 (2025).

58. G. Stecher, K. Tamura, S. Kumar, Molecular Evolutionary Genetics Analysis (MEGA) for macOS. Mol Biol Evol 37, 1237–1239 (2020).

59. S. Kumar, G. Stecher, M. Li, C. Knyaz, K. Tamura, MEGA X: Molecular Evolutionary Genetics Analysis across Computing Platforms. Mol Biol Evol 35, 1547–1549 (2018).

60. S. Q. Le, O. Gascuel, An improved general amino acid replacement matrix. Mol Biol Evol 25, 1307–1320 (2008).

61. D. N. Mastronarde, Automated electron microscope tomography using robust prediction of specimen movements. J Struct Biol 152, 36–51 (2005).

62. S. H. Scheres, RELION: implementation of a Bayesian approach to cryo-EM structure determination. J Struct Biol 180, 519–530 (2012).

63. J. Zivanov, T. Nakane, S. H. W. Scheres, Estimation of high-order aberrations and anisotropic magnification from cryo-EM data sets in RELION-3.1. IUCrJ 7, 253–267 (2020).

64. S. Q. Zheng et al., MotionCor2: anisotropic correction of beam-induced motion for improved cryo-electron microscopy. Nat Methods 14, 331–332 (2017).

65. A. Rohou, N. Grigorieff, CTFFIND4: Fast and accurate defocus estimation from electron micrographs. J Struct Biol 192, 216–221 (2015).

66. T. Bepler et al., Positive-unlabeled convolutional neural networks for particle picking in cryo-electron micrographs. Nat Methods 16, 1153–1160 (2019).

67. D. Kimanius, L. Dong, G. Sharov, T. Nakane, S. H. W. Scheres, New tools for automated cryo-EM single-particle analysis in RELION-4.0. Biochem J 478, 4169–4185 (2021).

68. T. D. Goddard et al., UCSF ChimeraX: Meeting modern challenges in visualization and analysis. Protein Sci 27, 14–25 (2018).

69. P. Emsley, K. Cowtan, Coot: model-building tools for molecular graphics. Acta Crystallogr D Biol Crystallogr 60, 2126–2132 (2004).

70. U. Litvin et al., Viro3D: a comprehensive database of virus protein structure predictions. bioRxiv, 2024.2012.2019.629443 (2024).

71. T. I. Croll, ISOLDE: a physically realistic environment for model building into low-resolution electron-density maps. Acta Crystallogr D Struct Biol 74, 519–530 (2018).

72. P. V. Afonine et al., Real-space refinement in PHENIX for cryo-EM and crystallography. Acta Crystallogr D Struct Biol 74, 531–544 (2018).

73. L. Holm, DALI and the persistence of protein shape. Protein Sci 29, 128–140 (2020).

74. L. Holm, Benchmarking fold detection by DaliLite v.5. Bioinformatics 35, 5326–5327 (2019).

75. L. Holm, Dali server: structural unification of protein families. Nucleic Acids Res 50, W210–W215 (2022).

76. H. Liu, A. Laiho, P. Toronen, L. Holm, 3-D substructure search by transitive closure in AlphaFold database. Protein Sci 34, e70169 (2025)

